# Talin-vinculin precomplex drives adhesion maturation by accelerated force transmission and vinculin recruitment

**DOI:** 10.1101/735183

**Authors:** Sangyoon J. Han, Evgenia V. Azarova, Austin J. Whitewood, Alexia Bachir, Edgar Guttierrez, Alex Groisman, Alan R. Horwitz, Benjamin T. Goult, Kevin M. Dean, Gaudenz Danuser

## Abstract

Talin, vinculin, and paxillin are mechanosensitive proteins that are recruited early to integrin-based nascent adhesions (NAs). Using machine learning, traction microscopy, single-particle-tracking, and fluorescence fluctuation analysis, we find that talin, vinculin, and paxillin are recruited in near-synchrony to NAs maturing to focal adhesions. After initial recruitment of all three proteins under minimal load, vinculin accumulates in these NAs at a ~5 fold higher rate than in non-maturing NAs and with faster growth in traction. We identify a domain in talin, R8, which exposes a vinculin-binding-site (VBS) without requiring load. Stabilizing this domain via mutation lowers load-free vinculin binding to talin, impairs maturation of NAs, and reduces the rate of additional vinculin recruitment. Taken together, our data show that talin’s concurrent localization with vinculin, before engagement with integrins, is essential for NA maturation, which entails traction-mediated unfolding of talin and exposure of additional VBSs triggering further vinculin binding.

## Introduction

Cell-matrix adhesions are macromolecular complexes that link the extracellular matrix (ECM), typically via integrin transmembrane receptors, to the actin cytoskeleton. Being both a force-transmitter and a force-sensor, cell-matrix adhesions are critical to cell morphogenesis and mechanosensation^1, 2^. Indeed, in response to ECM changes, adhesions undergo constant changes in morphology and motion that involve the recruitment and recycling of a large number of adhesion molecules. For example, nascent adhesions (NAs) emerge within the actin-dense cell lamellipodia and then slide in the direction opposite to the protrusion as a result of polymerization-driven flow of the actin network^1^. Many of these NAs, which are less than 0.5 μm long (and thus appear as diffraction or near-diffraction limited spots in fluorescence microscopy) are rapidly turned over and disassembled. A subset of NAs mature into longer focal complexes (FCs, >0.5 μm in length) and focal adhesions (FAs, >2 μm in length) at the lamellipodia-lamella interface^1, 3^. During this progression, NAs go through multiple decision processes regarding fate and morphology. Compared to the well-studied FAs, for which the interconnection between structure, signaling, and force transmission is largely understood^4–12^, much less is known about the molecular and mechanical factors that determine NA assembly, turnover, and maturation. In part, this is because until recently, it has not been technically feasible to measure whether individual NAs transduce traction forces. By combining high refractive-index and mechanically-tuned substrates that are compatible with total internal reflection microscopy^13^ and numerical methods for the computational reconstruction of cell-substrate traction at the single micron length-scale, we demonstrated that force transmission is essential for the stabilization and maturation of NAs^14^. However, it remained unknown which factors determine whether a NA begins to bear forces and thus continues to assemble.

One possible factor that may influence NA maturation is the stoichiometry of the earliest molecular components recruited to the site of its formation^15, 16^. In particular, the recruitment of talin, vinculin, and paxillin could play a critical role as they all are known to be mechanosensitive^17–24^. Talin is an integrin activator^25, 26^ that directly links integrins to the actin cytoskeleton^27^. Under force, the helix bundle domains in talin’s rod-like region unfold^19^, which both disrupts ligand binding and exposes cryptic binding sites for vinculin and other proteins^19, 28–31^. Vinculin, when bound to talin’s exposed binding sites, can indirectly strengthen the connection between actin and integrins by 1) forming a catch bond with F-actin^32, 33^, 2) establishing multivalent linkages between talin and F-actin filaments^28, 29, 34^, and 3) stabilizing talin’s unfolded state^35^. In this scenario, talin first binds to integrins and F-actin, is unfolded under the initial load, and serves as a scaffold for the recruitment of vinculin. Indeed, at the level of FAs, direct evidence for catch-bonds^36–38^, and the exposure of hidden binding sites under load^39, 40^, has established the idea of force-assisted adhesion growth. Further evidence for this model indicates that downregulation of actomyosin contractility reduces the recruitment of vinculin^24^ and other adhesion proteins^41^, as well as the association between talin and integrins^42^.

In contrast to a model of hierarchical FA growth and stabilization where talin arrives first and recruits vinculin, fluorescence fluctuation analyses^42^ and co-immunoprecipitation experiments^24^ have suggested that talin and vinculin might form a complex before talin associates with integrins. While talin-vinculin pre-association implies vinculin’s force-independent binding to talin, it is not clear whether this pre-association is required for NA assembly, and if so, whether pre-association affects the decision processes for NA maturation. Moreover, paxillin, a scaffolding protein that works in close relationship with focal adhesion kinase (FAK)^24, 43–45^, is thought to be recruited and stabilized by force at an early phase of NA assembly^5, 23, 46, 47^. However, vinculin’s recruitment and role in establishing tension across NAs remains poorly understood^48–50^.

Here, we investigated the integration of molecular recruitment and mechanical forces in determining the fate of NAs. Specifically, we combined high-resolution traction force microscopy with highly sensitive particle detection and tracking software to evaluate the time courses of force transmission and molecular composition of individual NAs. A comprehensive inventory of these traces revealed broad heterogeneity in NA behavior. Thus, we applied machine learning approaches to divide NAs into subgroups with distinct characteristics and identified that five subgroups are necessary to account for the different kinematic, kinetic and mechanical properties of NAs. By focusing on the NA subgroup that matures into stable FAs, we found that the formation of a talin-vinculin precomplex was mediated by talin’s R8 domain. These precomplexes reinforce the link between talin and actin, likely to allow the unfolding of talin and exposure of additional vinculin binding sites, which ultimately supports the transition of spontaneous molecular assemblies in nascent adhesions into stable macromolecular focal adhesions.

## Results

### Nine adhesion classes can be distinguished based on different kinetic and kinematic behaviors

To investigate the time courses of traction forces and protein recruitment in NAs, we performed two-channel time-lapse, total internal reflection fluorescence (TIRF) imaging of Chinese Hamster Ovary epithelial cells (ChoK1, Supplementary Fig. 1a). We chose this cell model because they form a stable footprint in close apposition to a flat substrate, which is ideal for TIRF microscopy of individual adhesions^16, 42, 51, 52^. In addition, these cells display a relatively slow progression of adhesion assembly, offering the opportunity to track the fate of NAs and to identify the requirements for maturation^46, 48, 49, 53^. ChoK1 cells were transiently transfected with low-levels of either talin-GFP, vinculin-GFP, or paxillin-GFP, and plated and allowed to spread and migrate on high-refractive index silicone gels. To measure traction forces, deformable silicone gels were densely coated with 40 nm fluorescent beads (2.2 ± 0.31 beads/μm^2^, 0.42 ± 0.17 μm bead-to-bead spacing, Supplementary Fig. 1b,c). After each experiment, cells were removed from the substrate and the beads imaged in the relaxed, undeformed configuration of the silicone gel, which permits the quantitative reconstruction of traction forces at sub-micron scales (Supplementary Fig. 1e,g,h)^14^. As expected, all traction force vectors pointed from the cell periphery to the cell center, independent of which adhesion protein was imaged (Fig. 1a-c). Likewise, regardless of which protein was ectopically expressed, ChoK1 cells spread and generated traction forces indistinguishably (Supplementary Fig. 2).

**Figure 1.**
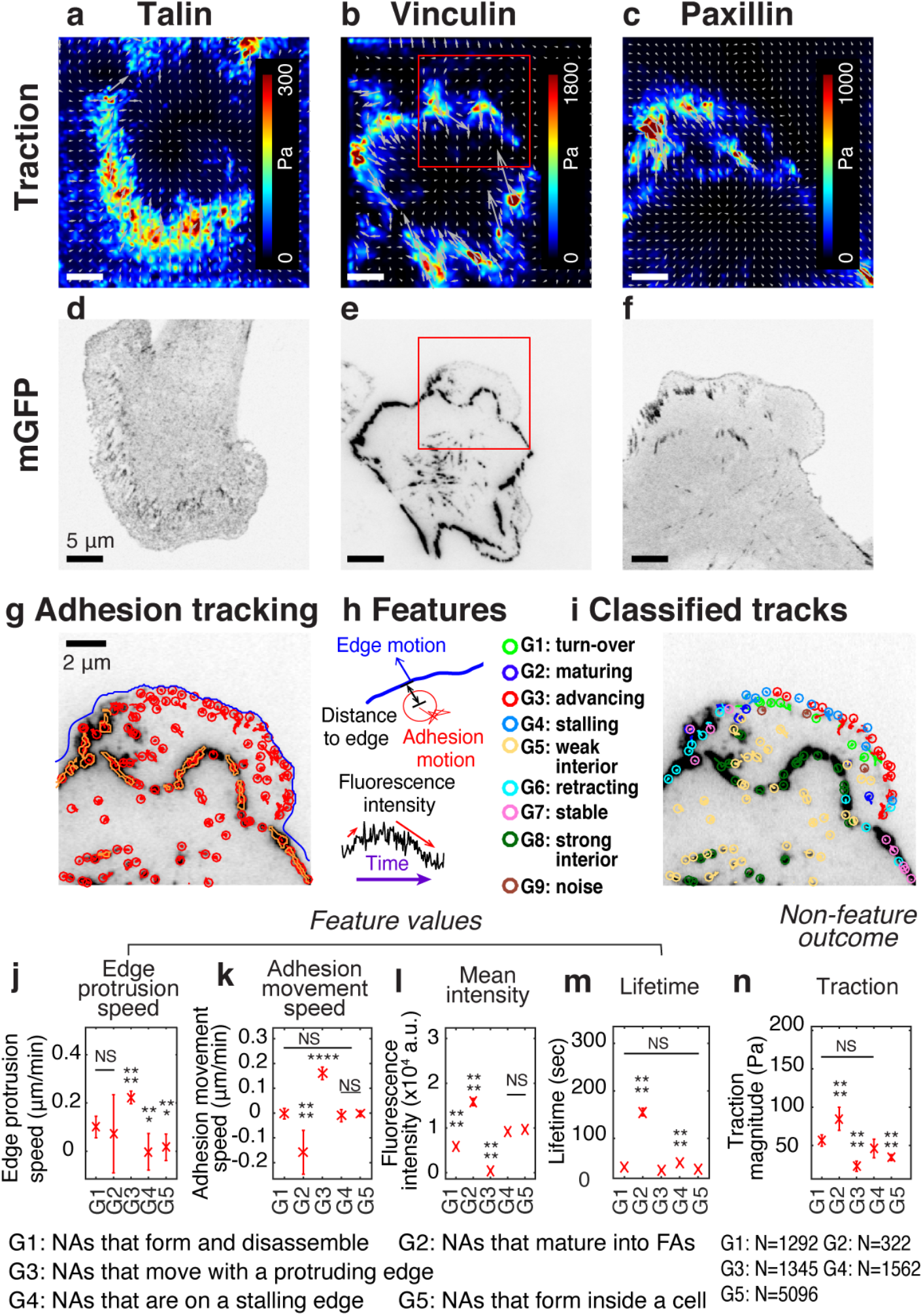
Experimental/computational pipeline to analyze heterogeneous adhesion dynamics in ChoK1 cells. (a – c) high-resolution traction maps co-imaged with mGFP-tagged adhesion protein, talin (d), vinculin (e), and paxillin (f). 5 kPa silicone gel coated with high density beads was used as a TFM substrate. (g) Trajectories of individual nascent and focal adhesions overlaid on a region of interest cropped from (e). Tracking is based on all detected point sources, (red circles). Big segmented focal contacts/adhesions (orange, closed freeform overlays) were used as additional information for feature selection. (h) Some of the key features used for supervised classification, tabulated in Table S1. (i) Classification of adhesion trajectories into nine different groups, overlaid on the adhesion image. Five different NA groups, three FA groups and one noise group were distinguished by the support vector machine classifier. (j-m) Comparison of feature values among the five NA groups, G1, G2, G3, G4 and G5: edge protrusion speed (j), adhesion movement speed, positive when sliding toward protruding edge (k), mean intensity (l), and lifetime (m), extracted from six vinculin-tagged cells. All features show a significant shift in value for at least one subgroup. (n) Average traction magnitude, read from traction map, at individual NA trajectories per each group. The number of samples per each group is shown in the lower right corner of the figure.

Fluorescently tagged adhesion proteins (Fig. 1d-f) were detected and tracked, and their intensity time courses extracted from the trajectories (Fig. 1g, Supplementary Fig. 1f). To account for the heterogeneity of adhesions, we collected 22 features from each trajectory (Fig. 1h, Supplementary Table 1) and classified the adhesions into nine groups (see Supplementary Table 2 for a summary of each group) with a supervised machine learning pipeline (see Methods, Software Availability). To generate training data, a human operator used a dedicated graphical user interface for labeling ~120 adhesion tracks (~10 tracks per group, out of ~10,000 tracks per movie). Based on these data we trained a support vector machine (SVM) classifier (validation accuracy: 70 – 80 %, Supplementary Fig. 3a). All features were inspected for redundancy and similarity (Supplementary Fig. 3b-c), and each group was distinct in terms of its Euclidian distance to the closest group in the feature space (Supplementary Fig. 3d). SVM-based classification of all trajectories that were excluded from the training data assigned each adhesion to one of nine different classes, G1, G2, ..., G9 (Fig. 1i). Five of the nine classes (G1-G5) identified NAs, three (G6-G8) identified FAs, and one group (G9) contained insignificant, noise-like trajectories (Supplementary Movie 1). The five NA classes significantly differed in terms of features such as “edge protrusion speed” (Fig. 1j), “adhesion movement speed” (Fig. 1k), “average fluorescence intensity” (Fig. 1l), and “lifetime” (Fig. 1m). For example, NAs classified into G3 form at the tip of the protruding edge and move forward with the protrusion. Of all NA classes, their fluorescence amplitude is lowest (Fig 1l). NAs classified into G2 form at the protruding edge but slide rearward relative to the substrate and mature to form larger FCs or FAs, and have the highest intensity and longest lifetime (Fig 1l-m). Indeed, all G2 NAs (~100%) mature to FCs, and ~32% mature into FAs (Supplementary Fig. 4). NAs classified into G1 also form at the protruding edge, but they are relatively stationary (Fig. 1k) and are characterized by a weak fluorescence intensity and a short lifetime (Fig. 1l-m).

### Nine adhesion classes exhibit distinct mechanical behaviors

Next, we tested the hypothesis that these spatially and kinetically distinct classes of NAs generated differential traction forces. Indeed, we found that the subgroup of maturing NAs, G2, showed the highest traction magnitude shortly after their initial assembly (Fig. 1n). This is consistent with previous findings that demonstrated the tension-mediated, myosin-dependent maturation of FAs^23, 46, 54^. Interestingly, NAs in G3 exhibited an insignificant amount of traction that did not increase throughout their lifetime (Fig. 1n, Supplementary Fig. 5), suggesting that this population of forward moving NAs might not interact with the oppositely directed F-actin flow. NAs in G1 had higher traction than those in G3, implicating that short-lived, non-maturing NAs can transmit significant amounts of traction forces, which is consistent with our previous findings^14^. Importantly, these trends were consistent regardless of which adhesion protein was used for tracking (talin, vinculin or paxillin; Supplementary Fig. 6). Furthermore, we observed shifts in the relative proportion of adhesion classes when cells were cultured on stiffer substrates (Supplementary Fig. 7). For example, large FAs (G8) were more abundant in ChoK1 cells cultured on 18 kPa substrates when compared to ChoK1 cells on 5 kPa substrates, which is a stereotypical output for stiffness sensing across adhesions ^9, 55^. At the same time, the population of most NAs, i.e., those in G1, G2 and G4, decreased for cells on 18 kPa relative to their 5 kPa counterparts. Altogether, these results confirm the reliability of the SVM classifier and suggest that kinetically unique NAs also show differences in terms of force transduction.

### Talin, vinculin and paxillin are recruited sequentially in non-maturing NAs, but concurrently in maturing NAs with traction development

To evaluate the relationship between molecular recruitment and traction force in NAs, we performed high-resolution traction force microscopy on cells labeled with either talin, vinculin, or paxillin, and processed these data using the aforementioned classifier (Supplementary Movies 1-9). We focused our analysis on the differences between NAs in G1 and G2 (Fig. 2), which are for simplicity henceforth referred to as non-maturing and maturing NAs, respectively. For all proteins evaluated, fluorescence intensity traces for non-maturing NAs had a lifetime of ~6-7 min, with clear rising and decaying phases (Fig. 2a-f, top). The associated traction traces exhibited intermittent rises and falls, but with an overall magnitude that was much smaller than the traction traces in maturing NAs (Fig. 2a-f, bottom). As expected, maturing NAs showed a steady increase in both fluorescence intensity and traction with a lifetime greater than 15 minutes (Fig. 2g-l). The fluorescence intensity and traction of individual non-maturing and maturing NAs reflected this stereotypical behavior, and so did the average behavior, i.e. a slight increase and fall for non-maturing NAs, and more steady increase for maturing NAs (Supplementary Fig. 8a-i). A further analysis with cohort plots, where traces of similar lifetime are grouped and separately displayed, revealed that average traces of many cohorts follow the stereotypical behavior (Supplementary Fig. 8j-o).

**Figure 2.**
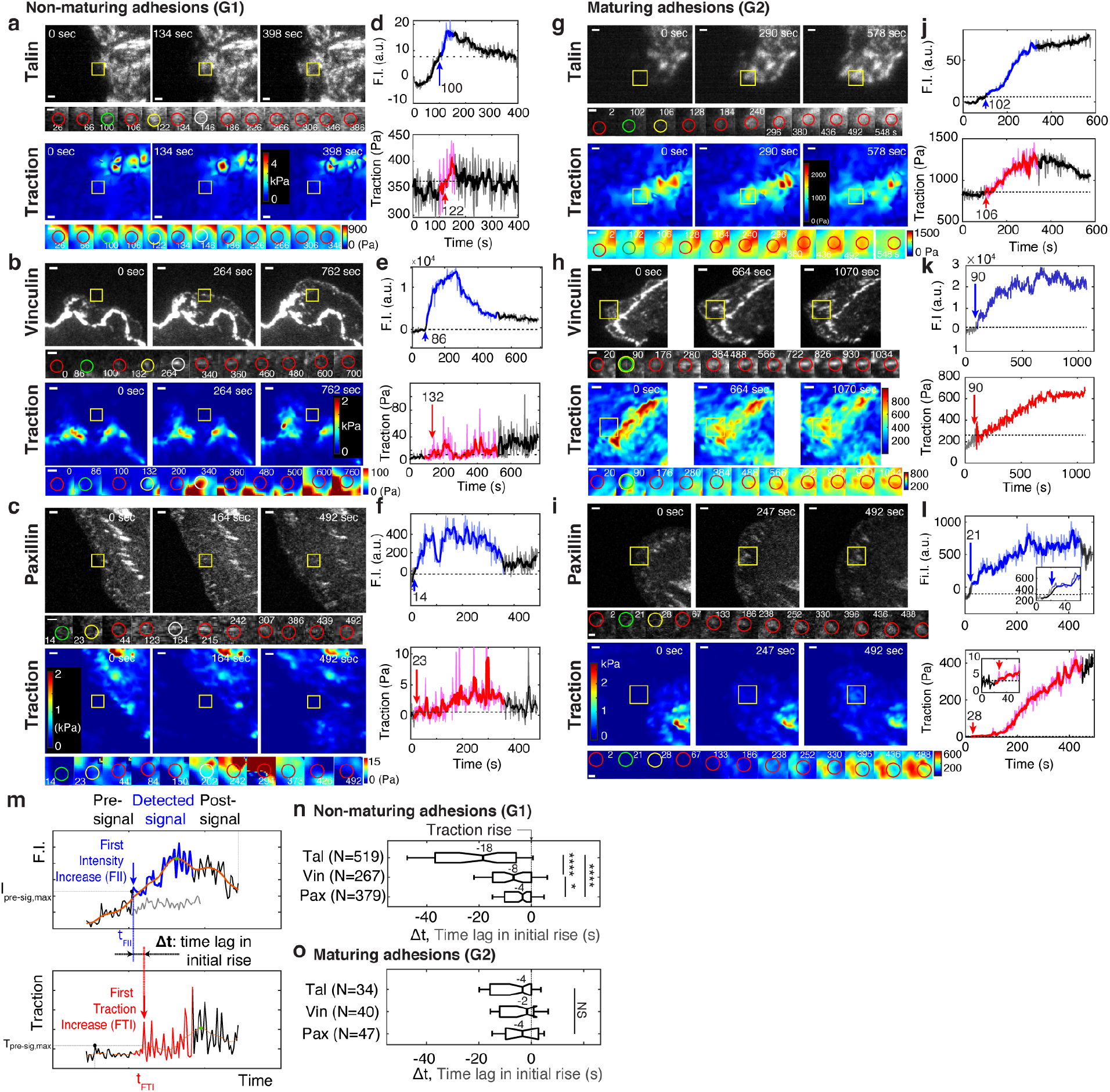
*Talin and vinculin in non-maturing NAs are recruited in a sequential manner before traction development whereas in maturing NAs they are recruited concurrently, along with paxillin, briefly before the initial traction rise.* (a-l) Representative traces of protein recruitment and traction generation at non-maturing (a,b,c), or maturing (g,h,i) NAs. Each panel contains three views at different time points approx. 2 – 4 apart of mGFP-tagged talin (a,g, top), vinculin (b,h, top), and paxillin (c,i, top) and associated traction maps (bottom). Yellow boxes indicate positions of example adhesions whose fluorescent signals and traction levels are shown in time lapse montages with finer resolution underneath. Scale bar: 1 μm. Green circle represents the time point of initial talin/vinculin/paxillin signal rise, yellow circles show the time point of initial traction rise, and white circles shows the time of the peak amplitude, while red circles show regular detections in between these events. Scale bar: 1 μm. (d-l) Traces of fluorescence intensity amplitude (top) and traction (bottom). Blue and red segments indicate periods of significant fluorescence intensity amplitude and of traction, respectively, illustrated as ‘detected signal’ in (m). The black segments indicate the remaining background-subtracted fluorescence intensity and traction levels outside the detected signal period, i.e., pre-signal and post-signal illustrated in (m). Whereas colored segments are read at positions moving with the adhesion center, pre- and post-signal traces are read at the first and last position of detected signal. An inset in (l) indicates that also in this trace the traction is gradually increasing. Blue and red arrows with the numbers on top mark the time points in seconds of the first intensity increase (FII) and the first traction increase (FTI), respectively, which are defined in (m). (m-o) Analysis of time-shifts between protein recruitment and FTI. (m) Traces of fluorescence intensity (top) and traction (bottom). Illustrated is the detection of the first significant value in both series. The grey signal represents the local background around the detected NA. Distinct distributions of time lags between FII and FTI in non-maturing (n) and maturing (o) NAs. Sample numbers, extracted from 6 cells for talin, 5 cells for vinculin and 4 cells for paxillin, are shown with each y-axis label. *: p<1×10^−2^, ****: p<1×10^−30^ by Mann-Whitney U test.

Next, we developed an event-based time-series analysis method that identifies the first time point of significant fluorescence and force increase, respectively, and then measures the time delay between the two (Fig. 2m). The blue and red arrows in Fig. 2d-f, j-l show, in two example traces, the time points identified statistically as the initial time point of intensity and traction increase, respectively. Using this approach, we first determined the fraction of NAs per group with a significant traction increase at any point throughout their lifetime (Supplementary Fig. 9). Interestingly, both non-maturing and maturing NAs showed such force increases, i.e. they were engaging and clutching against F-actin’s retrograde flow at one point with the substrate. In contrast, NAs classified as groups G3-5 exhibited lower fractions of force increase, suggesting that a very large number of adhesion protein complexes, detectable through either talin, vinculin, or paxillin recruitment, do not transduce measurable traction forces (e.g., less than 20 Pa). Indeed, these complexes could result from an incomplete molecular clutch and be engaged with the substrate but not actin, or vice versa.

To evaluate if maturing and non-maturing NAs were differentially assembled, we next analyzed protein recruitment using the initial traction force increase as a time fiduciary. In non-maturing NAs, talin, vinculin and paxillin were recruited ~18 sec, ~8 sec and ~4 sec before the onset of force transmission, respectively (Fig. 2n). In contrast, in maturing NAs, talin, vinculin and paxillin were recruited concurrently ~4 sec before the onset of force transmission (Fig. 2o). We also noted that the temporal distribution of talin recruitment was significantly wider in non-maturing NAs than in maturing NAs. These findings suggest that in maturing NAs talin localizes with vinculin prior to its association with integrins, which more efficiently leads to force transmission and maturation to a FC. These data also suggest that NAs that sense force at the earliest stages of assembly are more likely to successfully mature. In contrast, although non-maturing NAs eventually also support some lower level of force transmission (Fig. 1n), talin and vinculin assemble sequentially and remain under force-free conditions for a much longer duration of time.

### In maturing NAs, vinculin assembles faster than in non-maturing NAs, but talin and paxillin show no difference

The rod domain of talin contains 13 helical bundles, 9 of which include cryptic vinculin binding sites (VBSs) that are exposed after tension-mediated mechanical unfolding^7, 19, 30^. Thus, we hypothesized that the simultaneous talin-vinculin recruitment in maturing NAs could further accelerate vinculin binding compared to non-maturing NAs. To test this, we quantified the assembly rate of each protein by obtaining the slope of the fluorescence intensity over the first 10 sec after initial appearance (Fig. 3a). Interestingly, only vinculin showed a significant difference in the assembly rate between non-maturing vs. maturing vinculin complexes, while talin and paxillin showed no such differences (Fig. 3a). Thus, the concurrent arrival time of talin and vinculin in maturing NAs could prime talin to expose additional VBS domains, which in turn would reinforce further vinculin recruitment and adhesion maturation. We also quantified the traction force growth in those NAs with an expectation that there would be an immediate rise in force with faster vinculin binding. However, the traction force growth rate for the first 10 seconds showed no significant difference between non-maturing vs. maturing NAs (Fig. 3b). This observation could arise because the forces were below our limits of detection for traction, or because traction does not develop during the earliest stage of molecular recruitment. However, a difference was observed when the force growth rate was quantified over a longer period, i.e., two minutes (Fig. 3c), which is consistent with our previous observations^14^. These findings imply that increased vinculin recruitment in maturing NAs – because of the effective tension development across the talin/vinculin mediated linkage between integrin and F-actin – supports the rise of traction force by connecting the protein complex to more F-actin, with some time delay.

**Figure 3.**
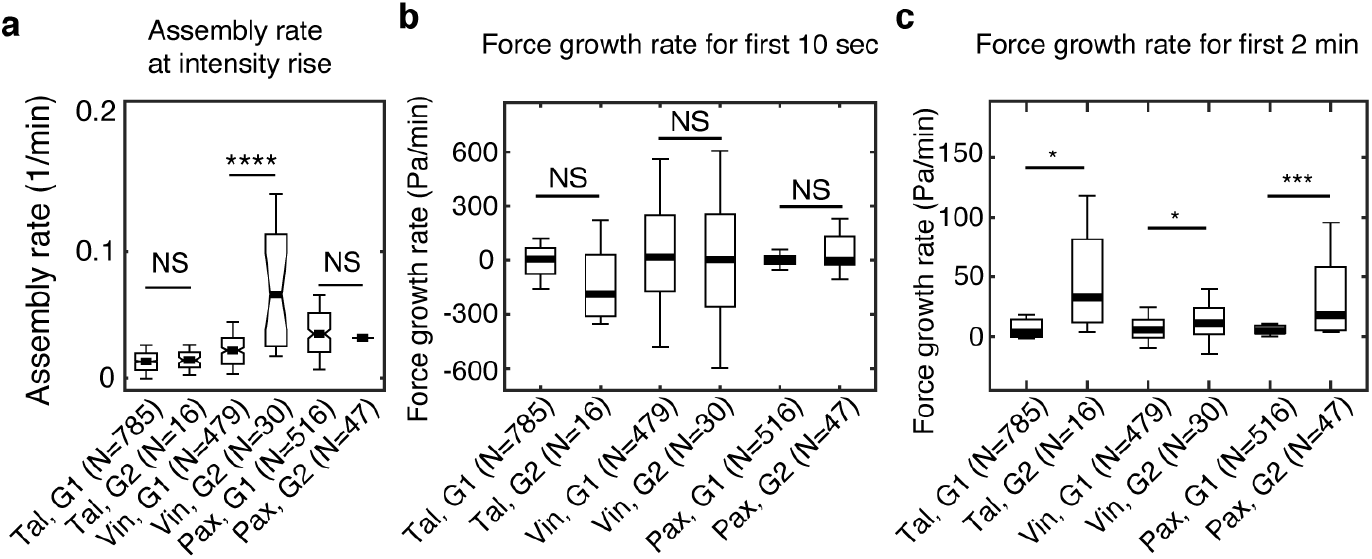
*Vinculin, but not talin and paxillin, is recruited significantly faster in maturing NAs than in non-maturing NAs.* (a) Assembly rate of talin, vinculin and paxillin, to G1 (non-maturing) or to G2 (maturing) NAs, quantified by the slope of fluorescence intensity over the initial 10 seconds after detection. (b) Traction growth rate at the NAs in (a) for the initial 10 seconds after detection. (c) Traction growth rate quantified over the first two minutes after detection. *: p<1×10^−2^, ***: p<1×10^−10^, ****: p<1×10^−30^ by Mann-Whitney U test

### Vinculin can bind to talin without force through a ‘threonine belt’ in talin R8 domain

Previous work established that under tension the R3 domain in talin unfolds first, as it contains a destabilized hydrophobic core due to the presence of a ‘threonine belt’. By mutating the threonine residues to isoleucines and valines (the so called “IVVI mutant”) it was possible to stabilize the core and prevent talin from unfolding, which significantly reduces the exposure of the two cryptic VBS ^30, 35, 55^. Moreover, we showed that the VBS in R8 was able to bind vinculin readily in the absence of force^56^. Like R3, R8 also contains a threonine belt, consisting of T1502, T1542 and T1562 (Fig. 4a)^29^. Thus, we hypothesized that a similar strategy, using a T1502V, T1542V, T1562V “R8vvv mutant”, could stabilize the R8 domain and reduce access to the VBS. To test this hypothesis, we made a “R7R8vvv” construct and compared its unfolding characteristics to wild-type R7R8 fragment (R7R8wt) using circular dichroism (CD; Fig. 4b). We included the R7 domain to improve the stability of the fragment and maintain R8 in its native conformation. In the R7R8wt the two domains unfolded cooperatively with a single unfolding step at a melting temperature (T_m_) of 55°C. In contrast, stabilization of the R8 domain in the R7R8vvv mutant resulted in the domains unfolding independently, with R7 unfolding at a similar temperature to the wt (T_m_ 56°C), but the melting temperature of the R8 domain increased from 56°C to 82°C. Strikingly, the two unfolding steps indicate that in the R7R8vvv mutant the R7 and R8 behave independently with regard to thermal stability. Together, these results show that the R7R8vvv mutant stabilizes R8.

**Figure 4.**
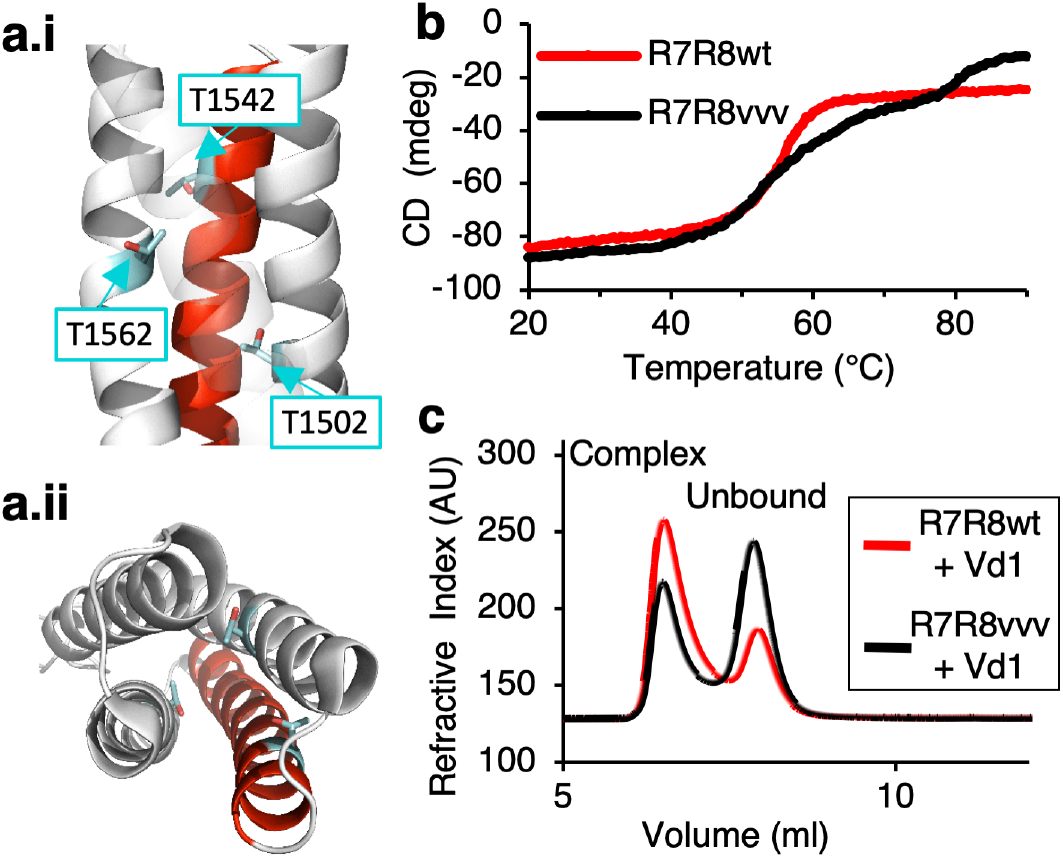
*Stabilizing the “threonine belt” in the R8 domain of talin inhibits talin-vinculin interactions under tension-free conditions*. (a) Cartoon representation of talin R7R8 (pdb id 2X0C) showing the ‘threonine belt’, comprised of residues T1502, T1542, and T1562, labeled and shown as sticks (cyan), the VBS helix is colored red. (a.i) side on view (N.B. helix 31 transparent), (a.ii) top down view. (b) Denaturation profiles for wildtype R7R8wt (red) and R7R8vvv (black) measured by monitoring the change in circular dichroism at 208 nm with increasing temperature. R7R8wt has a melting temperature of 55°C, whereas R7R8vvv unfolds in two steps, one (R7) with a melting temperature of 56°C and R8 unfolding at 82°C. (c) Chromatograms showing binding of talin R7R8 to the vinculin head (Vd1). R7R8wt (red) and R7R8vvv (black) binding to Vd1. Complex peaks and unbound peaks are indicated.

To test whether stabilization of R8 would affect its interaction with vinculin, we used analytical gel filtration to look at complex formation. Preincubation of R7R8wt or R7R8vvv with vinculin Vd1 showed that both constructs were able to form complexes. However, the R7R8vvv:Vd1 complex peak was substantially smaller than the wildtype:Vd1 peak, confirming that the R7R8vvv bound less Vd1 (Fig. 4c). Thus, by stabilizing the threonine belt with the R8 valine mutations, vinculin’s accessibility to the cryptic VBS was reduced. Interestingly, the R8 domain is also a Leucine-aspartic acid (LD) motif binding site, i.e. it binds to multiple proteins that act downstream of adhesion, including RIAM and DLC1^30, 31, 57^. Using a fluorescence polarization assay described previously^58^, we measured the binding affinities of R8 ligands RIAM TBS1 and DLC1 peptides with R7R8wt and R7R8vvv (Supplementary Fig. 10). Both peptides bound to the R7R8vvv with comparable affinities to the wildtype R7R8, confirming the R7R8vvv was still able to bind LD-motif proteins. Altogether, these biochemical characterizations suggested that the threonine belt in talin R8 is responsible for vinculin binding without force. The mutant also provided a tool for us to probe the functional implications of talin-vinculin precomplex formation on NA assembly and maturation *in vivo* without interfering with binding of other binding partners.

### Cells with R8vvv mutant talin show less maturing NAs and sparser and smaller FAs

To investigate whether talin’s ability to form a precomplex with vinculin promotes adhesion maturation, we introduced the R8vvv mutation into full-length talin1 and tagged it with mNeonGreen (“talin1 R8vvv-mNG”). For imaging, we knocked down endogenous talin1 expression with an shRNA, and rescued ChoK1 cells with shRNA-resistant forms of wild-type or R8vvv talin. The expression of talin1 R8vvv-mNG was slightly less than that of endogenous talin1 in wild-type ChoK1 cells, but more than the remaining endogenous talin1 expression in knockdown cells (Supplementary Fig. 11). Interestingly, cells expressing the talin1 R8vvv-mNG mutant contained many more NAs (Fig. 5a,b,f,g,k) and less and smaller FCs and FAs (Fig. 5a,b,f,g,l,m) than control cells expressing wild-type talin1. Furthermore, cells expressing the R8vvv variant of talin1 also showed less traction than cells rescued with wild-type talin1 (Fig. 5c,h,n). With less traction and more NAs, the edge protrusion and retraction velocity was also faster in cells expressing the talin1 R8vvv-mNG mutant (Fig. 5d,i). Moreover, a lower fraction of NAs and FCs in R8vvv mutant cells grew in size to FAs than NAs in cells with wild-type talin1 rescue (Fig. 5e,j,o,p). Together, these results demonstrate that talin R8vvv mutation restricts NAs from maturing into focal adhesions.

**Figure 5.**
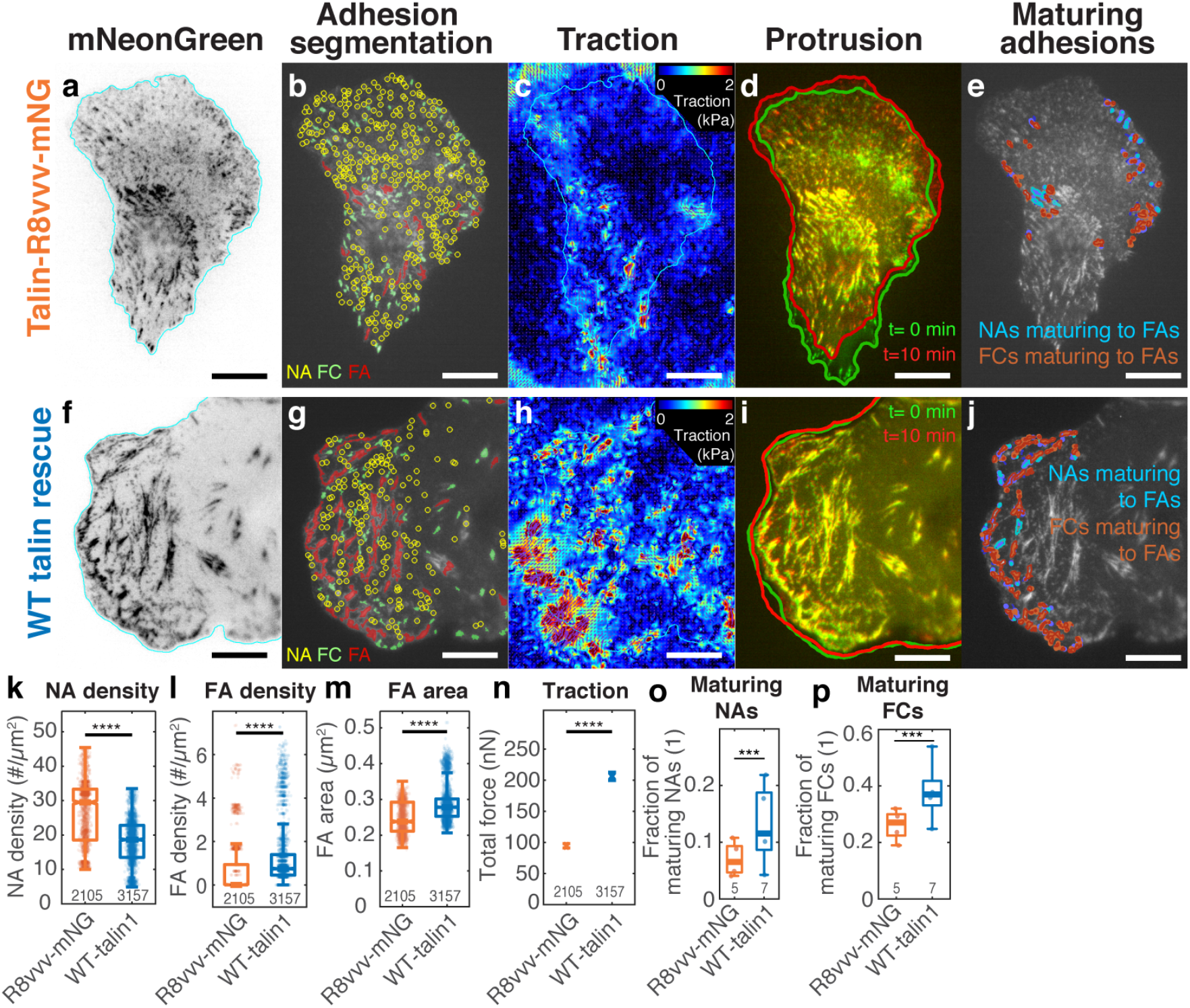
Expression of the talin1 R8vvv mutant in ChoK1 cells with endogenous talin1 knocked down results in the formation of denser NAs, but lesser and smaller FAs, lower traction, more active protrusions, and less maturing adhesions compared to wild-type talin. (a-j) Adhesion, traction and protrusion phenotype of a representative ChoK1 cell on 5 kPa substrate expressing (a-e) talin R8vvv mutant or (f-j) wild-type talin. (a and f) inverted talin-mNeonGreen images. (b and g) detection of NAs, FCs and FAs. (c and h) traction force maps. (d and i) snapshots of computer vision-extracted cell boundaries at 0 and 10 min of a movie. (e and j) overlay of NAs and FCs that mature to FAs. (k-p) Box plots of (k) NA density, (l) FA density, (m) FA area, (n) total traction integrated over cell area, (o) fraction of NAs maturing to FAs relative to all NAs, (p) and of the fraction of FCs maturing to FAs (relative to all FCs). Number of adhesions imaged are listed under each box plot. Number of independently imaged cells for talin1 R8vvv-mNG and WT talin1-mNG rescue were 5 and 7, respectively. Scale bar: 10 μm. ****: p<1×10-30 by Mann-Whitney U test

Noticeably, despite the reduced NA maturation, ChoK1 cells expressing the talin1 R8vvv mutant exhibited some large FAs (Fig. 5a,b and m). Accordingly, we asked if these FAs could reflect a population of NAs that mature under recruitment of residual endogenous talin1 (Supplementary Fig. 11), or perhaps talin2, which we did not specifically knockdown. Western Blot analysis showed that talin2 expression was minimal in ChoK1 cells compared to talin1 and knocking down talin1 did not induce compensatory expression of talin2 (Supplementary Fig. 12). The traction magnitude of ChoK1 talin1 knockdown cells was significantly lower than wild-type cells ectopically expressing talin1 or knockdown cells rescued with talin1 (Supplementary Fig. 13). Together, these data suggest that influence of a residual talin1 pool or of compensatory expression of talin2 on the behavior of ChoK1 cell with a talin1 knockdown and rescue by either R8vvv- or wild type talin 1 is minimal.

To ensure that the observed R8vvv NA maturation defect is independent of endogenous talin1 and talin2 expression, we lentivirally introduced the wild-type and the R8vvv variants of talin1 into talin1/2 null inner medullary collecting duct (IMCD) cells (see Materials and Methods).^59^ Clearly demonstrating the importance of talin for cell adhesion, talin1/2-null IMCD cells grew in suspension, and only adhered to the substrate when rescued with talin1 constructs. IMCD cells rescued with R8vvv-talin1 showed an adhesion phenotype that is very similar to ChoK1 rescued with R8vvv-talin1. Specifically, cells rescued with talin1 R8vvv had more NAs, smaller FCs and FAs, less traction, less NAs and FCs maturing to FAs than IMCD cells rescued with wild-type talin1 (Supplementary Fig. 14). Thus, we concluded that the remaining large FAs likely result from talin1-vinculin precomplexes, which still form in the presence of the R8vvv mutation, albeit to a lesser extent (See Figure 4C).

### R8vvv mutation does not affect talin recruitment but impedes traction growth rate

To investigate whether talin precomplex formation with vinculin affects talin recruitment itself, we compared the time of talin1 recruitment in R8vvv and wild type talin1 rescue cells for non-maturing (G1) and maturing (G2) NAs. Consistent with the data in Fig. 2, in both cases non-maturing NAs showed talin recruitment, on average, ~14 sec prior to the initial rise in traction (Fig. 6a,c,e,g,i), while maturing NAs showed an immediate talin recruitment (Fig. 6b,d,f,h,i). This indicates that the ability of talin to bind vinculin in the absence of the force does not affect talin recruitment. For both wild type and R8vvv mutant talin, the assembly rates were statistically indistinguishable between non-maturing and maturing NAs (Fig. 6j). The rate of traction development in NAs, however, was significantly affected in talin1 R8vvv-mNG mutant cells. Overall, the traction growth rate was reduced in R8vvv cells, both for non-maturing and maturing NAs (Fig. 6k). Moreover, whereas maturing NAs in WT talin1 rescue cells showed faster traction growth than non-maturing NAs, consistent with the data in Fig. 3c, maturing adhesions in talin1 R8vvv-mNG mutant cells exhibited an even slower force growth than non-maturing NAs (Fig. 6k). These results suggest that the R8vvv mutation does not change talin’s own recruitment rate, but interferes with force transduction in maturing NAs.

**Figure 6.**
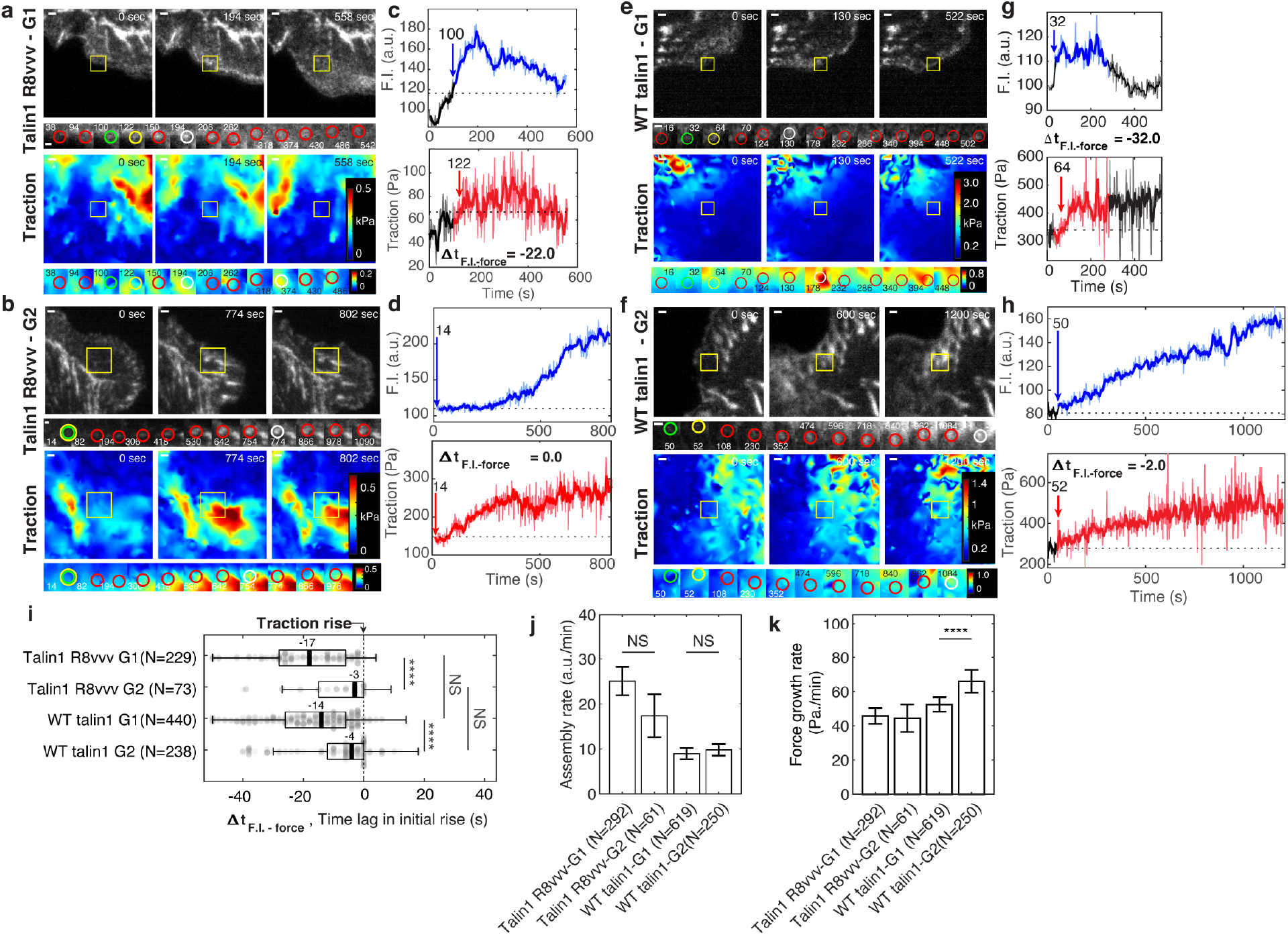
*Expression of talin1 R8vvv-mNG mutant does not change the recruitment timing of talin to NAs, but reduces the force growth rate in NAs.* (a-h) Representative talin (top) and traction force (bottom) images of talin1 R8vvv-mNG expressing cells (a-d) and WT talin-mNG rescue cells (e-h) within non-maturing (a,c,e,g) and maturing (b,d,f,h) NAs. (a,b,e,f) talin-mNG images (top) and traction images (bottom) of three different time points, i.e. at initial nucleation, at maximum fluorescence intensity, and at the end of the NA portion of the track. Yellow boxes indicate positions of example adhesions whose fluorescent signals and traction levels are shown in time lapse montages with finer resolution underneath. Green circles indicate the time points of initial talin signal rise, yellow circles show the time points of initial traction rise, and white circles show the time of the peak amplitude Red circles show normal default detections without special events. Scale bar: 1 μm. (c-d, g-h) Traces of talin-mNeonGreen fluorescence intensity (top) and traction (bottom). Phases of the traces with significant fluorescence above background are indicated in blue and red, respectively. The black time series outside the colored signal are the background-subtracted intensities read at the first or last position detected by the particle tracker. Blue and red arrows mark the time points of the first intensity increase and the first traction increase, respectively (i-k) Distributions of time lags of fluorescence intensity onset relative to traction onset (i), talin assembly rates (j), and traction growth rates (k) of non-maturing (G1) and maturing (G2) NAs in talin1 R8vvv-mNG mutant and WT talin1-mNG rescue cells. Time integration time for calculating slopes in j and k was 1 minute. *: p<1×10^−2^, **: p<1×10^−3^, ***: p<1×10^−10^, ****: p<1×10^−30^ by Mann-Whitney U test

### Differential vinculin recruitment between non-maturing vs. maturing NAs vanishes with talin1 R8vvv mutation

Vinculin recruitment to the NA is critical for both force growth and adhesion maturation (Fig. 3)^6^. To examine whether the assembly rate of vinculin is affected by talin’s ability to form a precomplex with vinculin, we performed two-channel imaging of vinculin-SnapTag-TMR-Star and wild type- or R8vvv talin-mNeonGreen (see Methods). As performed previously, we captured and analyzed time-series of each pair of talin-vinculin signals in non-maturing vs. maturing NAs (Fig. 7a-h) and quantified the vinculin assembly rate within 30 seconds after first detection (Fig. 7i). In talin1 R8vvv-mNG mutant cells, vinculin assembly rates were statistically indistinguishable between non-maturing (G1) and maturing (G2) NAs, whereas in wild type talin1-rescue cells vinculin rates were significantly higher in maturing NAs, consistent with the data acquired in control cells (Fig. 3a). This result suggests that early vinculin binding to talin R8 domain indeed contributes to faster recruitment of additional vinculin. The insignificant difference in vinculin recruitment in R8vvv mutant cells for non-maturing vs. maturing NAs might be related to the reverted traction growth rates between the two NA groups observed in these mutant cells (Fig. 6k). It is also worth noting that the vinculin signal in maturing NAs of cells with wild type talin-rescue tended to keep increasing while talin intensity was relatively flat (t=200~600 sec in Fig. 7h,d), suggesting that the number of exposed talin VBSs is increasing, and thus the number of bound vinculin proteins, over time under tension. The same trend was observed in talin1 R8vvv mutant cells (Fig. 7b,f), but the vinculin recruitment rate was again much less than those found in wild type talin1-rescue. Altogether, this data strongly suggests that vinculin recruitment is significantly reduced in the absence of a vinculin-talin precomplex.

**Figure 7.**
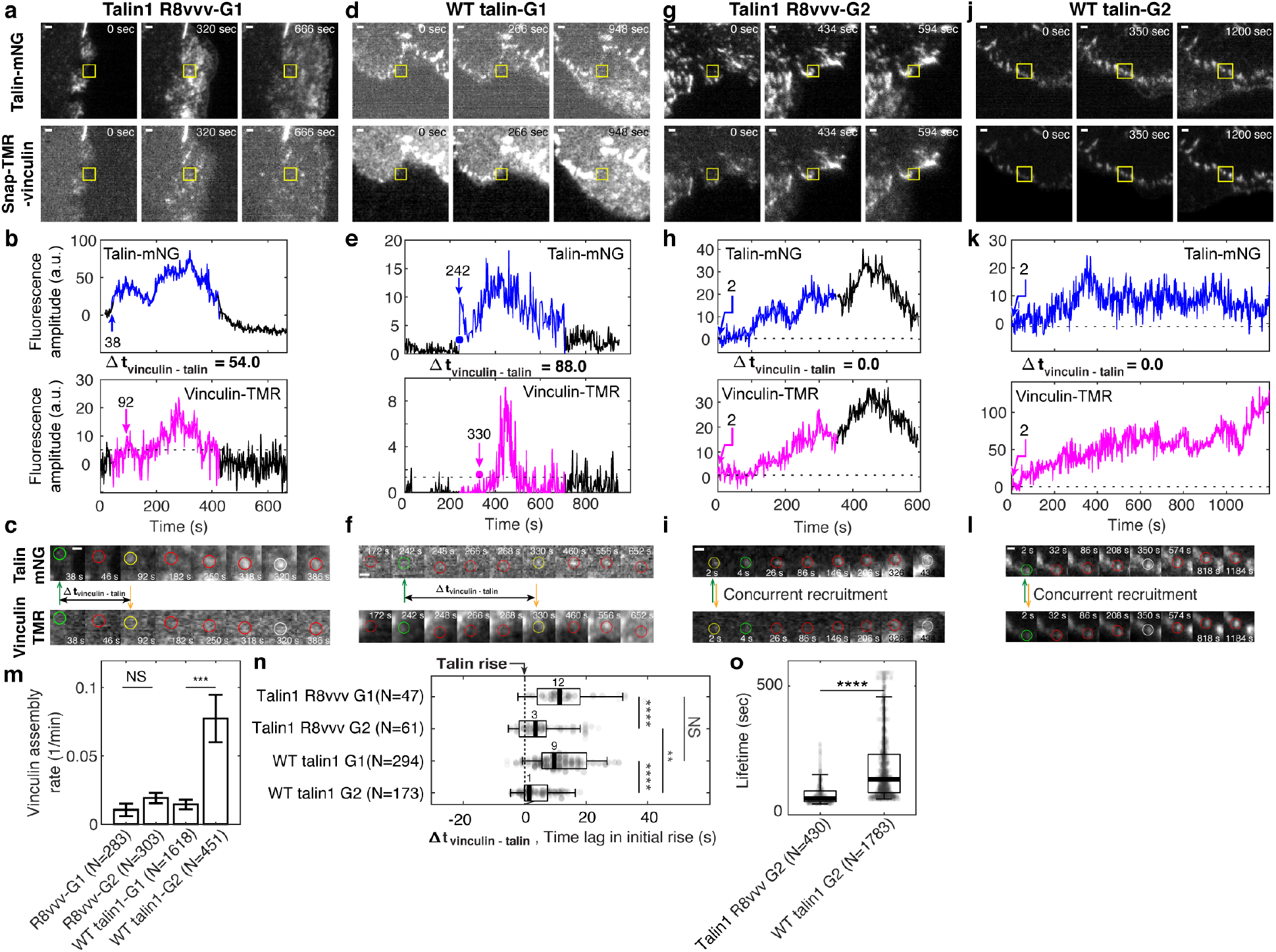
*Vinculin recruitment is reduced in talin1 R8vvv mutant cells.* (a,d,g,j) Representative two-channel time-lapse images of talin-mNeonGreen (top) and vinculin-SnapTag-TMR-Star (bottom) of G1 NA in a Talin1 R8vvv mutant cell (a), G1 NA in WT talin1 rescue cell (d), G2 adhesion in a Talin1 R8vvv mutant cell (g), and G2 adhesion in WT talin1 rescue cell (j). NAs of interest are indicated with a yellow box. Scale bar: 1 μm. (b-k) Time series of talin-mNeonGreen amplitude (top) and vinculin-SnapTag-TMR-Star amplitude (bottom) of G1 non-maturing (b,e) and G2 maturing (h,k) NAs in cells expressing the talin1 R8vvv mutant (b,h) and WT talin (e,k) constructs. Colored time periods (blue for talin, magenta for vinculin) indicate the phases where the adhesion is detected as a significant particle of robust trackability. The black time series outside the colored signal are the background-subtracted intensities read at the first or last position detected by the particle tracker. Blue and magenta arrows and the text around them indicate the time of talin and vinculin recruitment onset, respectively. (c,f,k,l) Time lapse montages of individual NAs shown in a, d, g, and j, respectively, overlaid with colored circles as detected centers of NAs of interest. Green circle represents the time point of initial talin signal rise, yellow the time point of initial vinculin signal onset, white the time of the peak amplitude, while red circles show normal default detections without special events. Talin and vinculin’s initial recruitments are indicated with green and yellow arrows to highlight the time delay occurring between talin and vinculin in G1 adhesions and the concurrent recruitment in G2 adhesions, regardless of R8vvv mutations. (m) Vinculin assembly rates at non-maturing and maturing NAs in R8vvv mutant and WT talin rescue cells, quantified by the first rate constant of vinculin-SnapTag-TMR-Star fluorescence intensity over the initial 30 seconds after the first detection in the talin-mNeonGreen channel. (n) Time delays of vinculin recruitment onset relative to talin recruitment onset of non-maturing vs. maturing NAs in talin1 R8vvv-mNG mutant and WT talin1 mNG cells. Vinculin recruitment onsets in non-maturing NAs are positive, i.e. vinculin recruitment starts after talin. In contrast, vinculin recruitment onsets in maturing NAs are nearly coincidental with talin. See the text for further description. (o) Lifetimes of maturing NAs classified in talin1 R8vvv mutant and WT talin1 mNG rescue cells. ****: p<1×10^−30^ by Mann-Whitney U test. The numbers of adhesions (N), extracted from 7 cells each for cells with talin1 R8vvv-mNG and WT talin1-mNG, are shown per each condition name at each panel.

### Simultaneous talin-vinculin imaging confirms vinculin’s recruitment after talin for non-maturing NAs and concurrent recruitment for maturing NAs

To confirm the recruitment order of talin and vinculin with respect to traction force development (Fig. 2), we quantified the time difference between the first significant increase in talin fluorescence intensity and the first significant increase in vinculin fluorescence intensity (blue and magenta arrows in Fig. 7a-h, j). For non-maturing NAs, both in talin1 R8vvv mutant and wild type talin1-rescue cells, vinculin was delayed to talin on average by ~10 seconds (Fig. 7j), consistent with the delay we inferred indirectly based on alignment of the fluorescent intensity increases with the first significant traction force increase (Fig. 2n). In maturing NAs, vinculin and talin recruitment coincided (Fig. 7j), also consistent with the indirect inference presented in Fig. 2n. This shows directly that the formation of a talin-vinculin precomplex indeed enhances the probability of NA maturation. In more detail, vinculin recruitment in maturing NAs of talin1 R8vvv mutant cells was ~4 seconds after talin recruitment, whereas vinculin recruitment in the wild type talin rescue condition preceded the talin recruitment by ~2 seconds (Fig. 7j). We interpret this difference as the result of the mutation in talin’s R8 domain, which reduces the ability of vinculin to bind talin prior to mechanical unfolding. Moreover, even though some maturing NAs eventually grow also in talin1 R8vvv mutant cells, the absence of efficient vinculin binding to the VBS in R8 propagates into an overall less efficient vinculin recruitment. In agreement with this interpretation, we found that the lifetimes of maturing NAs in the mutant cells were much shorter than those in cells with wild type-talin1 rescue (Fig. 7k). In sum, our data establishes that talin’s pre-association with vinculin via the talin R8 domain is critical for accelerated vinculin binding, which in turn contributes to the development of the level of force transmission required for NA maturation.

## Discussion

Our experiments show that the maturation of NAs depends on the formation and recruitment of a talin-vinculin precomplex. Previous models have inferred that tension across talin, which can establish direct bridges between integrins and actin filaments, is sufficient to unfold the molecule and expose several vinculin binding sites. These binding sites were thought to promote the recruitment of vinculin to further strengthen the link between the integrin-talin complex and actin^31, 63^. However, these models were derived primarily from observations in focal adhesions, i.e. at a late stage of the maturation process^6, 34^. Here, we exploit our ability to concurrently measure nanonewton-scale traction forces and molecular associations within individual NAs using total internal reflection microscopy on high-refractive index soft substrates^13, 14^. Our data suggests that the tension born by an individual talin bridge between integrin and actin filaments is insufficient to maximize the number of VBSs open for talin to form a stable link to F-actin. This further lowers the lifetimes of catch-bond-like molecular associations^33, 63, 64^ between talin and vinculin, vinculin and actin, and talin and actin, resulting in turnover of the NAs (Fig. 8, top). In contrast, pre-assembled talin-vinculin complexes immediately establish a strong link between integrin and F-actin, as indicated by the concurrent recruitment of talin and vinculin and traction force onset. The fast loading rate promotes an efficient unfolding of the talin rod domain, which exposes several additional VBSs for further recruitment of vinculin and strengthening of the talin/F-actin interaction. This results in robust increase of traction force transmission and stabilization of catch-bond-like molecular associations that contribute to the maturation of the NA (Fig. 8, bottom).

**Figure 8.**
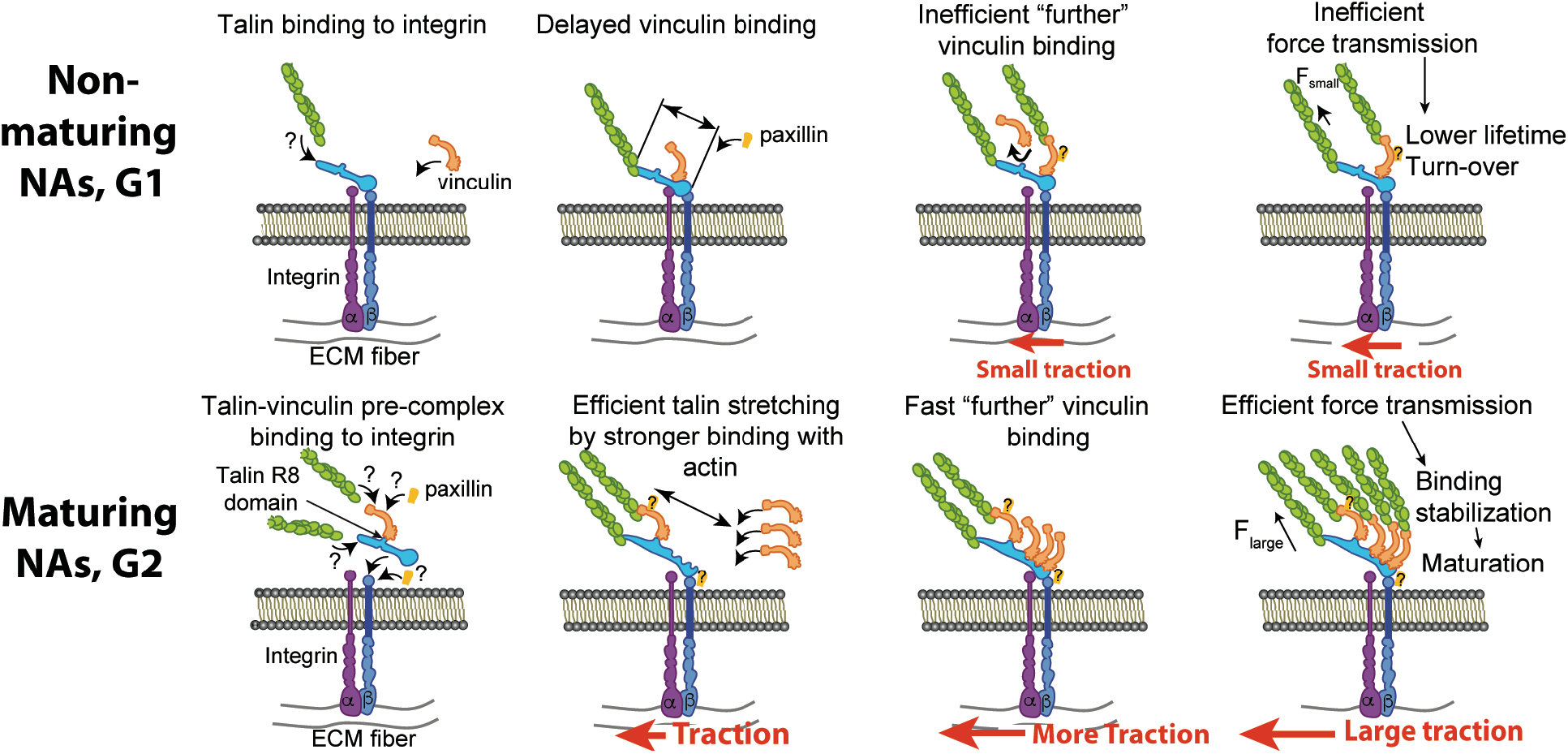
*A suggested mechanism of differential recruitment of talin and vinculin determining maturation of nascent adhesions.* (Top) For non-maturing NAs, talin binds to integrin before vinculin recruitment. Talin stretching might be limited to a shorter level, which limits the exposure of vinculin-binding-sites. Inefficient vinculin binding, in turn, limits the number of F-actin that can connect to the adhesion complex, allowing for only a low amount of tension across the complex. Insufficient loading level reduces the lifetime of catch-bond like associations between molecules, resulting in turnover of the NA complex. (Bottom) For maturing NAs, talin and vinculin form a precomplex before association with integrin. Upon precomplex recruitment to the NA traction force builds immediately. Talin might be stretched in a faster manner by pre-associated vinculin and talin’s own binding to F-actin accommodate faster, efficient recruitment of additional vinculin. High loading levels across the complex stabilizes molecular bonds, which facilitates the maturation of the NA. The sites for paxillin binding, e.g., to vinculin or β-integrin via FAK, are inferred from the literature ^22, 60–62^.

Our data also show that the colocalization of talin and vinculin is promoted by talin’s R8 domain, which contains a VBS that is exposed for vinculin recruitment without tension-mediated unfolding of talin. We generated a talin mutant with a more stable R8 domain that reduces the spontaneous association with vinculin. Cells expressing this mutant have a large fraction of NAs that cannot mature into FAs and transmit only low-level forces. Ultimately, our data suggests that maturing NAs are likely to be initiated by an R8-mediated talin-vinculin association. Intriguingly, complex formation prior to binding integrin receptors requires spontaneous encounters of mobile talin and vinculin at the plasma membrane or even within the cytosol. These are likely rare events, which may explain the surprising finding that the number of maturing (G2) NAs (3.5 ± 1.6 %, Mean ± Standard Error of the Mean, N=20 movies) is low compared to G1 NAs (28.8 ± 3.5 %, Mean ± S.E.M., N=20 movies) among all NAs.

Despite the general phenotype of less NA maturation, talin1-R8vvv-expressing cells also exhibited a small number of large FAs (Fig. 5j). These FAs are not due to residual endogenous talin1 or talin2 expression, as they are also evident in talin1/2 double knockout cells. Instead, these FAs in R8vvv-expressing cells likely result from one or more mechanisms. For example, 1) talin-R8vvv binding to vinculin is not abolished, only reduced (Fig. 4c), 2) talin-R8vvv’s association with integrins and F-actin could be quick enough to expose other VBSs, or perhaps 3) arise from one of the other NA classes (we focused largely on G1 and G2 here). Nonetheless, by performing three-color imaging of talin, vinculin, and traction forces (Fig. 7), we show that R8vvv cells do have maturing NAs, but less of them, with reduced lifetimes, and importantly, with impaired vinculin association rates (Fig. 7m). Thus, it appears that the second possibility might be less likely. Alternatively, we speculate that there may be a compensation effect at play where the fewer maturing NAs accumulate excess talin (Fig. 6j).

Our findings regarding the talin R8 domain offer an alternative perspective on talin-vinculin association, including our own report describing talin’s R3 domain as the weakest region that can unfold under tension^28, 34^. Likewise, there are recent results that seem conflicting with one another. Using truncated talin fragments, it would appear that the talin R4-R8 fragment does not bind the vinculin head domain, whereas the talin R1-R3 fragment does^65^. However, another recent study has shown that a talin lacking R2R3 is able to interact with both inactive and active vinculin^66^. Moreover, a few studies have supported an idea that R3 requires force, albeit small, for engagement with vinculin ^17, 28, 55^. First, a study with a talin tension sensor has shown that talin1’s engagement with the cytoskeleton must precede vinculin’s binding to the N-terminal VBS on talin1^17^. Second, it has been reported that IVVI mutant in talin R3 prevents mechanotransduction, as assessed by traction exertion and YAP nuclear localization^55^. Additionally, based on published data^30, 67^, the R3 domain is likely to be bound to the Rap1-interacting adaptor molecule (RIAM) prior to the force^68^. Thus, early, tension-independent interactions between talin and vinculin is more likely to mediated by a different site, for which our data suggests the talin R8 domain.

Our finding of a role for R8-mediated talin-vinculin complex formation in the earliest stages of adhesion assembly is also somewhat unexpected in view of the paradigm that both full-length talin and vinculin reside largely in an auto-inhibited conformation that prevents mutual interaction. Indeed, a recent study showed that full-length talin1 in a closed conformation does not form a complex with vinculin regardless of vinculin’s conformation^65^. In contrast, another study reported that only the ‘activated’ conformation of vinculin can form a complex with talin, or vice versa (i.e., an activated talin can form a complex with vinculin in its closed form)^66^. We speculate that both talin and vinculin are highly dynamic proteins that exist in an equilibrium that transitions between closed, open and intermediate states in living cells^65^, and the potential activation energy necessary for activation could easily arise from F-actin-driven forces. Additionally, although biochemically not yet confirmed, a talin-vinculin precomplex appears feasible as the crystal structure of talin shows an exposed R8 VBS. These considerations collectively align with observation that the occurrence of maturing G2 adhesions is less than the occurrence of non-maturing G1 adhesions. Thus, the conditions for efficient maturation, including the formation of a talin-vinculin complex without tension, are rarely fulfilled. Nonetheless, a rare but significant number of talin-vinculin precomplexes are sufficiently available to nucleate a population of G2 adhesions, which ultimately drives adhesion maturation.

In the case of non-maturing NAs, talin and vinculin were recruited significantly before our measurements could detect a significant traction onset. This finding implies that there are sub-populations of talin and vinculin that transduce little force. Additionally, talin is present for a longer time than vinculin before traction onset in non-maturing adhesions. While this is conceptually consistent with a previous finding that vinculin binding to talin requires talin’s actin binding for tension development^17^, the time lag between talin and vinculin recruitment suggests that talin’s sole engagement with F-actin without vinculin potentially impedes talin’s own role as an integrin activator ^69^ and promoter of integrin clustering ^70^. Additionally, before vinculin binding, talin may be bound to RIAM ^71^, which is replaced by vinculin only after the R2R3 domain unfolds ^30^.

How a talin-vinculin precomplex promotes faster tension development and talin unfolding in maturing adhesions remains to be determined. Potential mechanisms imply that 1) the complex is also pre-bound to F-actin through the vinculin tail (as vinculin bound to talin is almost certainly in an open conformation) ^22, 72^ and 2) that the talin-vinculin interactions via talin’s R8-domain do not interfere with talin’s direct binding to F-actin, thus accelerating talin’s actin-binding rate.

In maturing adhesions, paxillin is recruited concurrently with talin-vinculin precomplexes. Which molecular partners in NAs and binding sites within those molecules are responsible for paxillin’s concurrent binding needs further investigation. A previous study has shown that talin-vinculin interaction can facilitate efficient paxillin recruitment regardless of the paxillin-binding-site in vinculin’s tail domain^22^. This finding suggests that paxillin’s concurrent recruitment in maturing nascent adhesions is associated with the talin-vinculin precomplex but not necessarily via paxillin’s direct binding to vinculin^73^.

How non-maturing (G1) NAs switch to disassembly also necessitates further investigation. One potential scenario includes that talin is competitively bound by RIAM or DLC1 before it can associate with vinculin. Alternatively, vinculin binding to other VBSs within talin could interfere with force transduction and adhesion maturation (e.g., vinculin binding to R3, after partial unfolding by F-actin-mediated forces at ABS3). Whether vinculin’s binding to such VBSs, i.e., other than R8, interfere or synergize vinculin binding remains to be determined. However, our data suggests that when talin binds F-actin in G1 NAs – directly or indirectly via vinculin – the further recruitment of vinculin is much slower than that of G2 NAs where talin and vinculin arrive simultaneously in what we hypothesize is a precomplex.

Our data also indicates that the onset of traction force is accompanied by paxillin recruitment, regardless of the fate of the NA (Fig. 2). This suggests that, paxillin is recruited after vinculin, and this is particularly true in non-maturing NAs. Indeed, the tension dependency of paxillin recruitment is well-established^23^. Our data suggests that tension-dependent recruitment of paxillin is through vinculin, which is consistent with previous findings that suggest paxillin recruitment can be induced by vinculin^22^ as it binds to the tail-domain of vinculin^60^. Alternatively, paxillin has been reported to be recruited after focal adhesion kinase (FAK) in endothelial cells^61^. Given evidence that talin can also be recruited by FAK^62^, paxillin’s recruitment after talin and vinculin might be coincident with vinculin-paxillin binding mediated by FAK. In line with our measurements, a FRET-based tension sensor study recently showed that of the three molecules (paxillin, talin and vinculin), paxillin levels correlate strongest with traction force levels^74^. Altogether, our findings agree with previous evidence that paxillin levels are an accurate reporter of traction levels, but not NA nucleation.

In summary, our work establishes an unexpected role for a talin-vinculin precomplex as a mechanical prerequisite to the further recruitment of vinculin to talin, which is the foundation of adhesion maturation. While the possibility of talin-vinculin precomplexes has been discussed in previous studies^24, 42, 75^, their function has remained obscure until now. Here, we show that complex formation is an essential step in adhesion assembly. How this precomplex forms, and whether its formation is regulated by cellular signals, are two of the critical questions to be addressed in future studies.

## Materials and Methods

### Cell Culture

ChoK1 cells were cultured in Dulbecco’s Modified Eagle Medium (DMEM) with 4.5 g/L D-Glucose, L-Glutamine, and Sodium Pyruvate (Gibco, 11995-065) supplemented with 10% Fetal Bovine Serum (Equitech-Bio, Inc, SFBU30), 1% Anti-Anti (Gibco, 15240112), and 1% Non-Essential Amino Acids (Gibco, 11140076). For transfection, cells were plated in a 6-well plate at ~30% confluency and transfected the next day with 350 ng of fluorescent protein-, or SNAP-tagged adhesion marker, 1 μg of pBluescript (Stratagene) as non-specific DNA, 10 μL of Lipofectamine LTX (Gibco, 15338030), and 2 mL of reduced serum Opti-MEM (Gibco, 31985088) according to the manufacturer’s directions. Four hours after adding the DNA-lipid mixture to the cells, the media was replaced with full DMEM media. 24 hours later, cells were trypsinized, and enriched with flow cytometry for low-level GFP-positive cells. Of this pool, 50,000 cells were seeded on fibronectin-coated (see below) traction-force microscopy substrates in pH 7.4 HyQ-CCM1 media (GE Lifesciences, SH30058.03), supplemented with 1.2 g/L of sodium bicarbonate and 25 mM HEPES. <u>mGFP-talin1</u> (provided by N. Bate and D. Critchley), <u>paxillin-eGFP</u> (provided by I. Schneider), and <u>mGFP-vinculin</u> (provided by M. Humphries) were used for adhesion-TFM two-channel experiments.

### Knock-out and Knock-down Experiments

For knock-out experiments, Murine Inner Medullary Collecting Duct (IMCD) Talin1/2 knockout cells (the kind gift of Dr. Roy Zent, Vanderbilt University^59^) were grown in DMEM/F-12 Dulbecco’s Modified Eagle Medium/Nutrient Mixture F-12 with L-Glutamine, 15 mM HEPES (Gibco, 11330-032) 10% Fetal Bovine Serum (Equitech-Bio, Inc, SFBU30), and 1% Anti-Anti (Gibco, 15240112) under standard cell culture conditions (5% CO2, 37°C) and passaged regularly at ~70% confluency. Owing to the complete knockout of Talin1/2, these cells are nonadherent and propagate in suspension. To rescue Talin1, IMCD cells were separately transduced using lentivirus with mNeonGreen-tagged wild-type and R8 mutant Talin1 (T1502V, T1542V, and T1562V) at low-levels using a truncated cytomegalovirus promoter (pLVX-CMV-100, https://www.addgene.org/110718/). To generate lentivirus, human embryonic kidney cells were transiently transfected with transfer vector (pLVX-CMV-100-mNG-Talin-18/pLVX-CMV-100-mNG-Talin-18-R8), viral packaging (psPAX2, https://www.addgene.org/12260/) vector, and viral envelope (pMD2.G, https://www.addgene.org/12259/) vector, at a 1:1:1 ratio (5 mg each) using 250 ml of Opti-MEM (Gibco, 31985088) and 45 ml (1 mg/ml) of polyethyleneimine. The final mixture was incubated for 15 minutes at room temperature and then transferred dropwise to human embryonic kidney cells. After 48 hours, the viral supernatant was gently removed from the human embryonic kidney cells, filtered with low protein binding 0.45 mm syringe filters, and added to IMCD Talin1/2 knockout cells at ~70% confluency with 12 micrograms/mL of polybrene. 24 hours later the media was replaced with full DMEM/F-12 media, and changes in the adhesion behavior were observed 48 hours post-transduction. Approximately one-week post-infection, non-adherent cells were washed from the dish, and the remaining knockout cells were trypsinized and evaluated for traction force microscopy.

For knock-down experiments, a previously validated shRNA hairpin against talin (GGAAAGCTTTGGACTACTA), located on the N-terminus of talin1, was stably introduced into ChoK1 cells with a pLVX-shRNA1 lentiviral system (Clontech) and selected for with 5 μg/mL of puromycin. Western blot analysis indicated decreased levels of talin expression (Supplementary Figure 11). For rescue experiments, talin1 from mouse (Mus musculus) was subcloned into the pCDNA3.1(+) mammalian expression vector (ThermoFisher Scientific, V79020). To make the reconstitution vectors insensitive to the shRNA, silent mutations were introduced into the corresponding shRNA target sequence. The new sequence of the reconstitution vectors is now GGAA<u>G</u>GC<u>CC</u>T<u>A</u>GACTACTA. This silent mutation was introduced into mNeonGreen talin1 pcDNA3.1 vector, which was used for wild-type reconstitution. The pCDNA3.1-mNG-Talin1-shRNA vector was mutated for talin1 R8vvv mutant with mutation sites at T1502V, T1542V, and T1562V. The R8 mutations (T1502V, T1542V, and T1562V, according to mouse numbering), alter the stability of the talin R8 domain. For vinculin imaging, mNeonGreen (Allele Biotechnology) was replaced with SNAP-Tag using seamless cloning, and labeled with SNAP-Cell TMR-Star (New England Biolabs, S9105S) or SNAP-Cell 647-SiR (New England Biolabs, S9102S) according to manufacturer’s recommendations. All protein-coding regions of expression constructs were verified with traditional primer walking and Sanger sequencing.

### Expression of Recombinant Polypeptides

For in vitro analyses, murine vinculin Vd1 (residues 1–258), murine talin R7R8wt (residues 1357-1653) and R7R8vvv (residues 1357-1653; T1502V, T1542V and T1562V) were cloned into a pET151 vector (Invitrogen) and expressed in E.coli BL21(DE3) cells cultured in LB. Standard nickel-affinity chromatography was used to purify the His-tagged recombinant proteins as described previously^58^. The proteins were further purified using anion exchange chromatography following cleavage of the 6xHis-tag with TEV protease. Protein concentrations were determined using their respective extinction coefficients at 280 nm.

### Circular Dichroism (CD)

Spectroscopy was performed using a JASCO J-715 spectropolarimeter equipped with a PTC-423S temperature control unit. Denaturation profiles were measured from 20-80°C at 0.2°C intervals by monitoring the unfolding of α-helices at 208 nm. 0.1 mg/mL of protein was dissolved in phosphate buffered saline (PBS). Measurements were made in a quartz cell of 0.1 cm path length.

### Fluorescence Polarization Assays

To determine if other binding partners of talin R8 domain except for vinculin can still bind to R7R8vvv fragment, the relative binding affinities were measured using an in vitro fluorescence polarization assay. The R8 interacting, LD-motif containing peptides from DLC1 and RIAM, i.e., DLC1_465-489-C (IFPELDDILYHVKGMQRIVNQWSEK-C) and RIAM_6-30-C (DIDQMFSTLLGEMDLLTQSLGVDT-C), were coupled to a thiol-reactive fluorescein dye via the terminal cysteine. Peptides with a C-terminal cysteine were synthesized by GLBiochem (China). Stock solutions (i.e., peptide + fluorescein) were made in phosphate-buffered saline (PBS; 137 mM NaCl, 27 mM KCl, 100 mM Na2HPO4, 18 mM KH2PO4, pH 7.4), 1 mM TCEP and 0.05% Triton X-100. Excess dye was removed using a PD-10 desalting column (GE Healthcare, Chicago, IL, USA). Titrations were performed in PBS using a constant 1 μM concentration of fluorescein-coupled peptide with increasing concentration of R7R8 fragment (either wild type or vvv mutant); final volume 100 μM in a black 96-well plate. Fluorescence polarization (FP) measurements, in which the binding between the two polypeptides results in an increase in the fluorescence polarization signal, were recorded on a BMGLabTech CLARIOstar plate reader at room temperature and analyzed using GraphPad Prism. K_d_ values were calculated with nonlinear curve fitting using a one-site total binding model.

### Analytical Gel Filtration

Gel filtration was performed using a Superdex-75 size exclusion chromatography column (GE Healthcare) at a flow rate of 0.7 mL/min at room temperature in 50 mM Tris pH 7.5, 150 mM NaCl, 2 mM DTT. A sample of 100 μL consisting of 100 μM of each protein was incubated at a 1:1 ratio at 25°C for 10 minutes. The elution was monitored by a Malvern Viscotek SEC-MALS-9 (Malvern Panalytical, Malvern, UK).

### Western Blot

Cells were transfected under identical conditions as they were for imaging experiments but with a 10 cm dish and sorted with a flow cytometer (FACS Aria II SORP) for low expression. Cells were lysed by adding 2x laemmli + 10% b-ME, vortexing, and heating at 95°C for 10 minutes. Protein concentration was measured, and the same amount was loaded for each lane. The gel was semi-dry transferred with a turbo blot, and then incubated overnight in 5% milk in tris-buffered saline with 0.1% Tween 20 (TBST) at 4 degrees. Protein was visualized with an anti-talin antibody at 1:1000 and the loading control was visualized with anti-b-actin at 1:5000, each in 0.5% milk/TBST overnight at 4°C. Gels were then rinsed with TBST, and probed with IgG:horseradish peroxidase in 0.5% milk/TBST at 4°C for 1 hour and then at room temperature for another 30 minutes. Gels were rinsed three times for 20 minutes in TBST and then detected with enhanced chemiluminescence.

### TFM Substrate Preparation

All silicone substrates had a diameter of 35 mm, a stiffness of 5 kPa (with the exception of Supplementary Figure 7), were embedded with 580/605 or 640/647 (λ_EX_/λ_EM_) 40 nm-diameter beads, and were compatible with total internal reflection fluorescence illumination. Substrates were coated with fibronectin (Sigma Aldridge, F1141) the same day as imaging experiments were conducted by mixing 20 μL of a 10 mg/mL 1-ethyl-3-(3-dimethylaminopropyl) carbodiimide hydrochloride (EDC) solution, 30 μL of a 5 mg/mL fibronectin solution, and 2 mL of Ca^2+^ and Mg^2+^ containing Dulbecco’s Phosphate Buffered Saline (DPBS, Gibco, 14040117) for 30 minutes at room temperature. Thereafter, the substrate was rinsed 2 times with DPBS, and incubated with 2 mL of 0.1% (w/v) bovine serum albumin in DPBS for another 30 minutes at room temperature, and rinsed several times with PBS prior to seeding with 50,000 transiently transfected cells.

### Live-cell TIRF Imaging for TFM and Adhesion Proteins

After being seeded, the cells were allowed to adhere to the substrate for ~1 hour prior to imaging. This was one of the more effective ways to make sure that cells had active protrusions. Cells were imaged with a DeltaVision OMX SR (General Electric) equipped with ring-TIRF, which mitigates laser coherence effects and provides a more homogeneous illumination field. This microscope is equipped with a 60x, NA=1.49, objective, and 3 sCMOS cameras, configured at a 95 MHz readout speed to further decrease readout noise. The acquired images were in 1024×1024 pixel format with an effective pixel size of 80 nm. Imaging was performed at 37°C, 5% carbon dioxide, and 70% humidity. To maintain the optimal focus, laser-based identification of the bottom of the substrate was performed prior to image acquisition, with a maximum number of iterations set to 10. Laser powers were decreased as much as possible, and the integration time set at 200 milliseconds, to avoid phototoxicity. At the back pupil of the illumination objective, the laser power for both 488 and 568 nm lasers was ~44 μW. Imaging was performed at a frequency of 1 Hz for 5-10 minutes, and deviations between the alignment for each camera were corrected in a post-processing step that provides sub-pixel accuracy. After imaging, cells were removed from the substrate with a 30% bleach solution, and the beads on the relaxed gel substrate were imaged for each cell position. Rapid imaging was necessary to mitigate swelling effects in the silicone substrate and to resolve traction forces in nascent adhesions.

### TFM Force Reconstruction

Bead images of the deformed gel – acquired when a cell was on the substrate – and a ‘reference bead image’ of the relaxed gel acquired after cell removal - were processed for traction reconstruction as described previously^14^. Briefly, the bead images of the deformed gel were compared with the reference image using particle image velocimetry. A template size of 17 to 21 pixels, and a maximum displacement of 10 to 80 pixels, depending on the bead density and overall deformation, were used for cross-correlation-based tracking of the individual beads. The displacement field, after outlier removal, was used for traction field estimation over an area of interest. The area of interest on the reference bead image was meshed with square grids of the same width, which depends on the average area per bead. The forward map, which defines the expected deformation of the gel at all bead locations given a unit force at a particular mesh of the force grid, was created by solving Boussinesq Eq. under the assumption of infinite gel depth. This forward map was then used to solve the inverse problem, i.e. given the measured field of bead displacements, the underlying traction field is determined. The solution to this inverse problem is ill-conditioned in that small perturbations in the measured displacement field can yield strong variation in the reconstructed traction field. To circumvent this problem, the traction field was estimated subject to L1-norm regularization. As discussed in detail in ^14^, L1-norm regularization preserved the sparsity and magnitude of the estimated traction field. Also as discussed and validated in ^14^, the application of L1-norm regularization over the L2-norm regularization most traction force microscopy studies employ is essential to resolve force variation at the length scale of the distances between individual nascent adhesions. The level of regularization is determined by a control parameter. We chose the parameter based on L-curve analysis, which guaranteed a fully automated and unbiased estimate of the traction field ^14^. Strain energy, which represents the mechanical work a cell has put into the gel, was quantified as 1/2 × displacement × traction, integrated over a segmented cell area. The unit of this integral is femto-Joule.

### Adhesion Segmentation, Detection and Tracking

Focal adhesions (FAs) and diffraction-limited nascent adhesions (NAs) were detected and segmented as previously described^14^. Briefly, FAs from images of either labelled paxillin, talin, or vinculin were segmented with a threshold determined by a combination of Otsu’s and Rosin’s algorithms after image pre-processing with noise removal and background subtraction. Segmented areas larger than 0.2 μm^2^ were considered for focal contacts (FCs) or FAs, based on the criteria described by Gardel et al.^3^. Individual segmentations were assessed for the area and the length, which is estimated by the length of major axis in an ellipse that fit in each FA segmentation. FA density was calculated as the number of all segmentations divided by the cell area. Nascent adhesions were detected using the point source detection described in ^76^. Briefly, fluorescence images were filtered using the Laplacian of Gaussian filter and then local maxima were detected. Each local maximum was then fitted with an isotropic Gaussian function (standard deviation: 2.1 pixels, i.e. ~180 nm) and outliers were removed using a goodness of fit test (*p* = 0.05). NA density was defined as the number of NAs divided by the entire cell area.

### Adhesion Classification

From the adhesion tracks, features 1-9 in Supplementary Table 1 were captured from the individual fluorescence intensity traces, and features 10-21 in Supplementary Table 1 from the corresponding spatial properties, some in reference to the position and movement of the proximal cell edge and to the overlap with segmentations of focal adhesions and focal complexes. A total of 9 class outputs were used to train the adhesion tracks. We believe that nine classes are a minimum number that represents a heterogeneity of the cell-ECM adhesions in a cell because the five were dedicated to classify NAs. The three classes assigned for FA classes can be viewed as insufficient in terms of representing a whole FA population and can be expanded further into a larger number of classes, but they were sufficient in this study because NAs were its main scope.

The classification was accomplished using a cubic support vector machine (SVM), which was proven to be the most accurate among linear classifiers. The classifier was evolved in a human-in-the-loop fashion, i.e. the user iteratively adjusted machine-generated classifications. The initial training data was labeled with qualitative criteria described in Supplementary Table 2. To facilitate the labeling process, an automatic, filtering-based, labeling was also employed (see Supplementary Table 3).

Both manual labeling and automatic labeling have advantages and drawbacks in terms of classification accuracy: while the manual labeling is less consistent due to subjectivity and human error, the automatic labeling has deficiencies in terms of incompleteness of the filtering criteria in capturing all essential properties of different adhesion classes. To overcome these drawbacks, both methods were employed in a way that the automatic labeling was performed first, and then manual labeling was added for insufficiently-labeled classes. During the manual labeling, adhesion classifications were immediately updated and presented to the user to allow class reassignments of selected adhesions. The labeling process was regarded to be completed once at least 10 adhesions were labeled for each class. To remove classification bias due to potential imbalance in the number of labels across the classes, the minority classes were oversampled, and the majority classes were undersampled, based on the mean number of labels ^77^. After training a classifier on one movie, for a new movie, another iteration of automatic- and-manual labeling was executed to update the classifier, which was applied to predict the classes of adhesions in the movie. Separate classifiers were built for talin-, vinculin-, and paxillin-tagged adhesions.

### Statistical Methods

Processed data such as traction, assembly rate, time delays, lifetimes, adhesion densities and areas in different conditions were compared using Mann-Whitney non-parametric test since all individual distributions do not follow a normal distribution. Testing for normal distribution was done by one-sample Kolmogorov-Smirnov test using a ‘kstest’ function in Matlab.

### Software Availability

A GUI-based Matlab software is shared via GitHub at https://github.com/HanLab-BME-MTU/focalAdhesionPackage.git.

## Acknowledgement

We would like to thank Roy Zent (Vanderbilt) and Olga Martha Viquez (Vanderbilt) for providing IMCD talin 1/2 double KO cells. We also thank Joseph Chi and Dana Reed for preparing DNA constructs and assisting with western blots, and Assaf Zaritsky (Ben Gurion University) for providing helpful comments about machine learning. This work was funded by the following grants: NIH R15GM135806 (S.J.H.) NIH F32GM117793 (K.M.D.), NIH P01GM098412 (A.G., A.R.H., and G.D.), NIH R35GM136428 (G.D.), Biotechnology and Biological Sciences Research Council grant BB/N007336/1 (B.T.G.) and a Human Frontier Science Program grant RGP00001/2016 (B.T.G.).

## Author contribution

S.J.H., A.R.H and G.D. conceived the project. K.M.D. and E.V.A. performed imaging experiments with talin constructs and its variations (e.g. R8vvv mutant). K.M.D. created ChoK1 cells with talin1 shRNA and talin1 R8vvv expression. E.V.A. cultured IMCD talin1/2 KO cells and expressed WT talin1 and talin1 R8vvv vectors. A.B. performed imaging experiments for WT GFP-tagged vinculin and paxillin. S.J.H designed the experiments and performed TFM reconstruction, nascent adhesion analysis and machine learning from the images. S.J.H. and G.D. wrote the manuscript and S.J.H. made the figures. B.T.G. provided suggestions for talin structure and mutation. A.J.W. and B.T.G. performed biochemical experiments and analyses. Silicone substrates were provided by A.G. and E.G. All authors reviewed and provided feedback on the manuscript.

## Competing Interests

The authors declare no competing interests.

## Notes

### Competing Interest Statement

The authors have declared no competing interest.

### Summary of Updates

The main update in this revision was a new experimental validation where we used talin1/2 double knockout cells (IMCD cells, a gift from Dr. Roy Zent, Vanderbilt), expressed mNG-talin1 R8vvv mutant, and compared adhesion phenotypes to WT talin-GFP rescue. The result, that the R8vvv-expressing cells show less nascent adhesion maturation with smaller traction transmission than WT talin1 rescue, was consistent with our previous experiment with ChoK1 cells with talin1 shRNA silencing and re-expressing the mutant. Both results demonstrate that a potential force-free complex formation of talin and vinculin before actively linking integrin and F-actin is critical for the fate determination of nascent adhesions to mature or not.

