## Supplemental Information for "Talin-vinculin precomplex drives adhesion maturation by accelerated force transmission and vinculin recruitment"

Supplementary Table 1 Features used for classification of adhesions

| | Feature name | Definition | Targets | Range | Unit |
|---|---|---|---|---|---|
| 1 | Max Intensity | Maximum intensity over adhesion lifetime, normalized by the max intensity | high for G2, low for G1 | 0-1 | 1 |
| 2 | Mean intensity | Mean intensity over adhesion lifetime, normalized by the max intensity | high for G2, G6, G7 and G8, low for G1, G3, G4 and G5 | 0-1 | 1 |
| 3 | Initial intensity | Fluorescent intensity of the track at the start of the track, normalized by the max intensity | High for G7, low for both G1 and G2 | 0-1 | 1 |
| 4 | Relative intensity decay | (Max intensity – intensity at the end of the track) / Max intensity | G1: big G2: small | 0-1 | 1 |
| 5 | Lifetime | Adhesion lifetime | G9: short<br>G2, G7, G8: long | 1-600 | Sec |
| 6 | Rise time | Time from adhesion initiation to time of max intensity<br>Time at max intensity – time at the beginning of the track | High for G2 | 1-600 | Sec |
| 7 | Overall intensity slope | Slope of intensity over entire adhesion life time | Positive for G1 and G2 | | a.u./sec |
| 8 | Early intensity slope | Slope of intensity during initial one minute | Positive for G1 and G2 | | a.u./sec |
| 9 | Late intensity slope | Slope of intensity during late one minute | G1 < G2 | -1 to 1 | a.u./sec |
| 10 | Initial adhesion motion | Adhesion motion in the direction of protrusion for the first minute of lifetime. | Positive for G3 and G4 | | µm |
| 11 | Terminal adhesion motion | Adhesion motion in the direction of protrusion for the last minute of lifetime. | Negative for G4 and G6 | | µm |
| 12 | Max edge fluctuation | Maximum fluctuation of local edge protrusion during adhesion lifetime | High for G1, G2, G3 | | µm |
| 13 | Distance to edge at initiation | Closest distance of adhesion to the cell edge at initiation | Low for G1, G2, G3, G4, G7, high for G5 and G8 | | µm |
| 14 | Distance to edge at termination | Closest distance of adhesion to the cell edge at termination of adhesion's life | Low for G3 and G4, medium for G1 and G2, high for G5 and G8 | | µm |
| 15 | Change in distance to edge during lifetime | Difference in closest distance of adhesion to the cell edge for entire adhesion lifetime | Low for G7, negative for G6 | | µm |
| 16 | Variance of edge fluctuation | Standard deviation of edge fluctuation of local edge protrusion during adhesion lifetime | Zero for G9 (adhesions near image boundary) | | µm |
| 17 | Focal adhesion area | Segmented area for focal complexes (>0.5 µm in length) and focal adhesions (>2 µm in length) | Positive for G2 and G8 | 0.1 to >10 | $\mu m^2$ |
| 18 | Adhesion status at termination | The time of the last 'FA' status compared to the time at termination of adhesion lifetime, normalized with the adhesion lifetime. E.g. 0 when the adhesion is ending as a FA status at the end of the movie. | Near zero for G2 and G8. Intermediate value (e.g. 0.3-0.7) for G1, G3 and G4. | 0 to 1 | 1 |
| 19 | Time period as NA before FC or FA | Time period of the adhesion as 'NA' status before the first 'FC' or 'FA' status. | Near zero for G7 and G8. At least 20 seconds for G2. | 0 to the time of a movie | sec |
| 20 | Edge protrusion speed | An average speed of the edge closest to the adhesion in the projected direction outward normal to the edge for adhesion lifetime. | Positive for G1, G2, G3, negative for G6 | -1 to 1 | µm/min |
| 21 | Adhesion movement speed | A speed of the adhesion in the projected direction outward normal to the edge for adhesion lifetime. | positive for G3, slightly negative for G1 and G2, highly negative for G6 | -1 to 1 | µm/min |
| 22 | Texture homogeneity | Out of area of 10x10 pixel around x-y position, gray-level co-occurrence matrix is calculated with an offset of | High for true focal adhesions G2 and G8, low for nascent adhesions (G1, G3, G4) | 0 to 1 | 1 |

| | | | |
|---|---|---|---|
| | | (0,1), then homogeneity property is calculated as $\sum_{i,j} \frac{p(i,j)}{1+\lvert i-j \rvert}$ where p is the co-occurrence matrix, i and j are row and column indices, respectively. | even when the intensity is high. |

*Supplementary Table 2. Nine adhesion groups that are defined heuristically.*

| Class | Short name | Qualitative description |
|---|---|---|
| G1 | NAs turn-over | NAs that form at the edge but stays at their positions or slide rearward as the edge protrude forward and go on turn-over. |
| G2 | NAs maturing | NAs that form at the edge but stays at their positions or slide rearward as the edge protrude forward and *mature* into FCs and FAs |
| G3 | NAs moving along protruding edge | NAs that form at the edge and move forward with protruding edge. |
| G4 | NAs at stalling edge | NAs that form at an initially protruding edge and stay at the edge which becomes stalling. |
| G5 | NAs inside | NAs with low fluorescence intensity at the cell interior |
| G6 | FAs retracting | FAs or FCs at the retracting edge |
| G7 | FAs stable at the edge | Stable adhesions (FAs) at the static cell edge |
| G8 | FAs stable inside | FAs with high fluorescence intensity and long lifetime at the cell interior |
| G9 | noise or very transient | Insignificant tracks with short life time (< 5 sec), low intensity or large random movement |

*Supplementary Table 3. Automatic labeling criteria. The sign '-' means that the corresponding criteria was not used for automatic labeling.*

| | Adhesion Classes | G1 | G2 | G3 | G4 | G5 | G6 | G7 | G8 | G9 |
|---|---|---|---|---|---|---|---|---|---|---|
| Automatic labeling criteria | Edge velocity | Positive | Positive | Positive | Negative | Small | - | Positive | - | |
| | Relative distance to edge | Increasing | Increasing | Constantly small | - | Constantly small | Large | - | Large | Large |
| | Starting location | At the edge | At the edge | At the edge | - | - | - | At the edge | - | - |
| | Initial trend of amplitude | Rising | Rising | - | - | - | - | - | - | - |
| | Ending trend of amplitude | Decaying | Non-decaying | - | - | - | - | - | - | - |
| | Relative time of maximum intensity | Early in the lifetime | Late in the lifetime | - | - | - | - | - | - | - |
| | Overlapping with FA segmentation | Yes, but only up to small area | Yes, especially at the end | No | - | - | - | - | - | - |
| | Mean amplitude | High enough | High enough | - | - | High | Low | Low | High | Low |
| | Initial amplitude | Low enough | Low enough | - | - | - | - | - | - | - |
| | Lifetime with 'NA' status (not overlapping with segmentation) | At least 10 sec | At least 10 sec | - | - | - | - | - | - | - |
| | Image texture homogeneity | - | High | - | - | - | Low | - | - | - |
| | Adhesion velocity | - | - | Positive | Negative | - | - | - | - | - |
| | Slope of amplitude | - | - | - | Negative | - | - | - | - | - |
| | Lifetime | - | - | - | - | Long | Short | - | Long | At least twice of G6 |
| | Edge velocity at last 2 minutes of an adhesion's lifetime | - | - | Positive | - | - | - | Near zero | - | - |

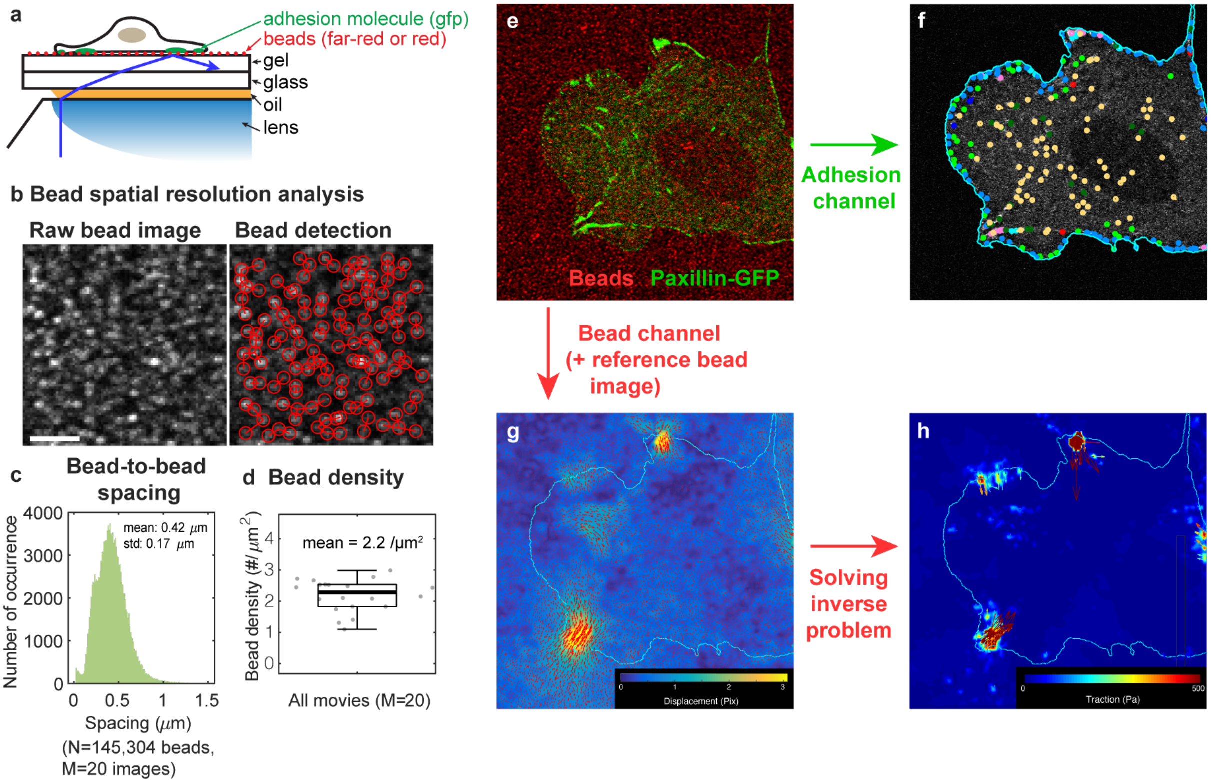

*Supplementary Figure 1. Simultaneous TFM-adhesion experimental approach. (a) A schematic of the TFM-adhesion imaging using TIRF. Here, cells adhere to a fibronectin-coated, high-refractive index, mechanically defined substrate that is compatible with TIRF. 40 nm beads on top of the substrate allow cell-mediated traction forces to be visualized. (b) Bead spatial resolution analysis. Beads on the silicone gel surface were detected using Gaussian-mixture model, and the inter-bead distance was calculated using KD-Tree algorithm (0.42 ± 0.17 μm, mean ± standard deviation). The bead density calculated from 20 bead images was 2.2 beads/$\mu m^2$ on average. (e-h) Analysis procedure for simultaneous TFM-adhesion imaging. (e) Overlay of adhesion channel (green) and bead channel (red). (f) Adhesions are analyzed for detection, tracking, and classification as described in the main manuscript and in Fig. 1. (g) Bead displacement fields are calculated by comparing the substrate state before and after removing the cells from the substrate. (h) bead channel with a reference image taken after the cells were removed from the substrate is analyzed for bead displacement. (f) Traction fields are obtained from the displacements solving the inverse problem as in Han et al. 2015.*

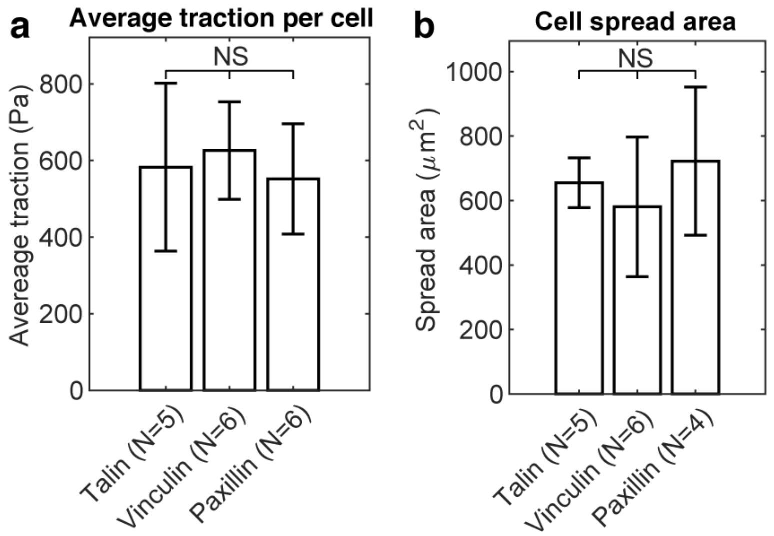

*Supplementary Figure 2. Overall average traction per cell and cell spreading area did not change with expression of talin-GFP, vinculin-GFP, or paxillin-GFP. (a) Bar plot of average traction quantified per cell. (b) Bar plot of cell spread area.*

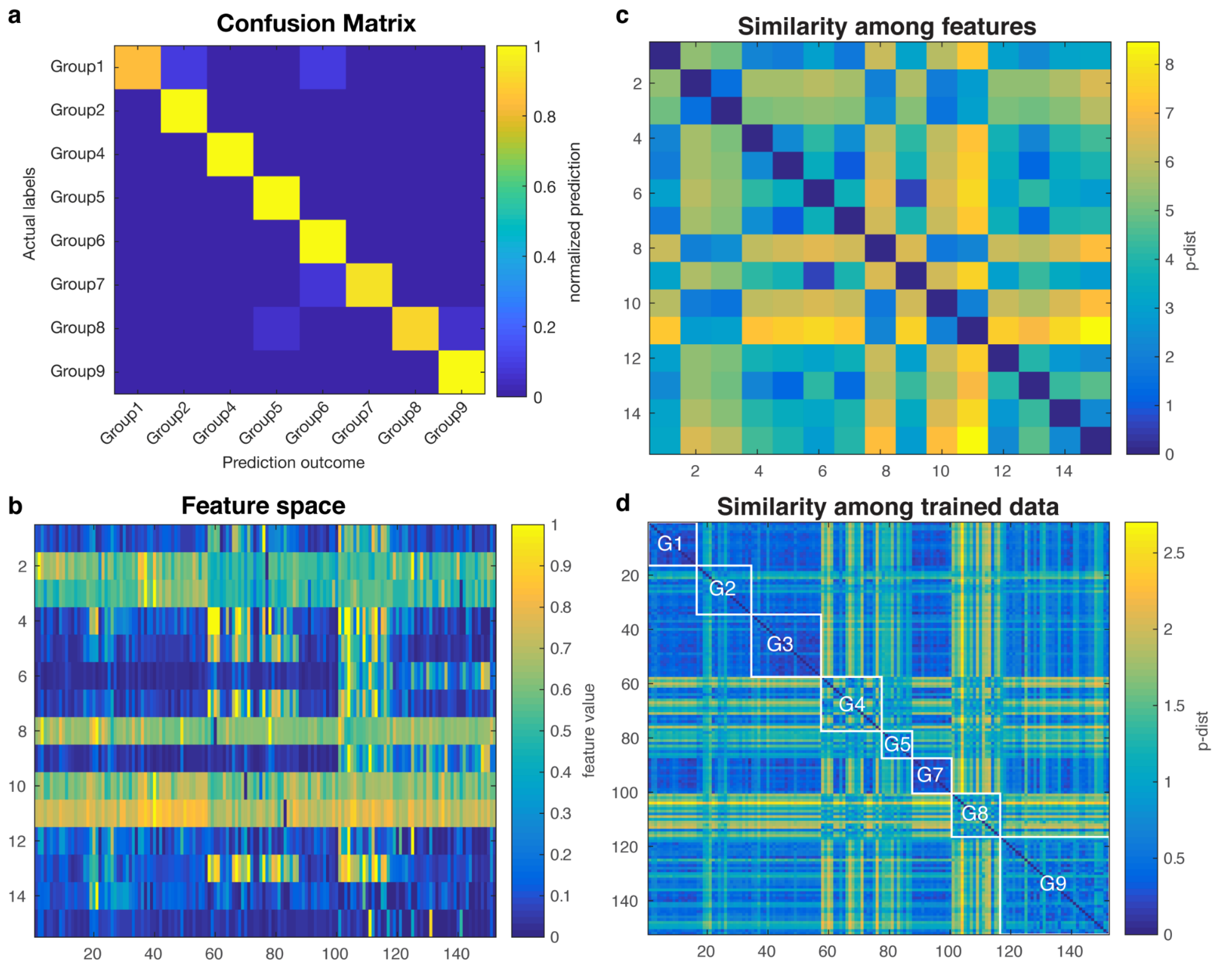

*Supplementary Figure 3. Validation of SVM-based machine learning. (a) confusion matrix among 9 different adhesion groups. (b) Feature space. (c) Similarity among features. (d) Similarity among trained data. White lined boxes represent similarity within each group.*

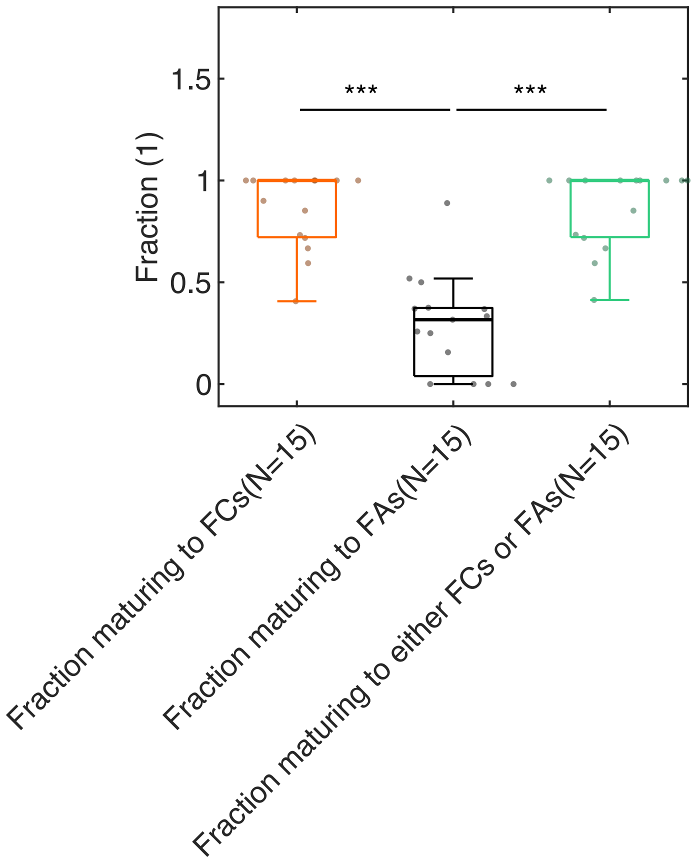

*Supplementary Figure 4. Boxplot of the fraction of G2 NAs that mature into FCs, FAs, or either FCs and FAs. N is the number of movies, which includes data obtained from cells expressing tagged variants of talin, vinculin, or paxillin. In total 10,028 G2 NAs were analyzed. Importantly, owing to the limited duration of imaging (10 minutes), this quantification is an underestimation since some (e.g., some G2 NAs do not progress to FC or FA because the movie concludes before this can occur).*

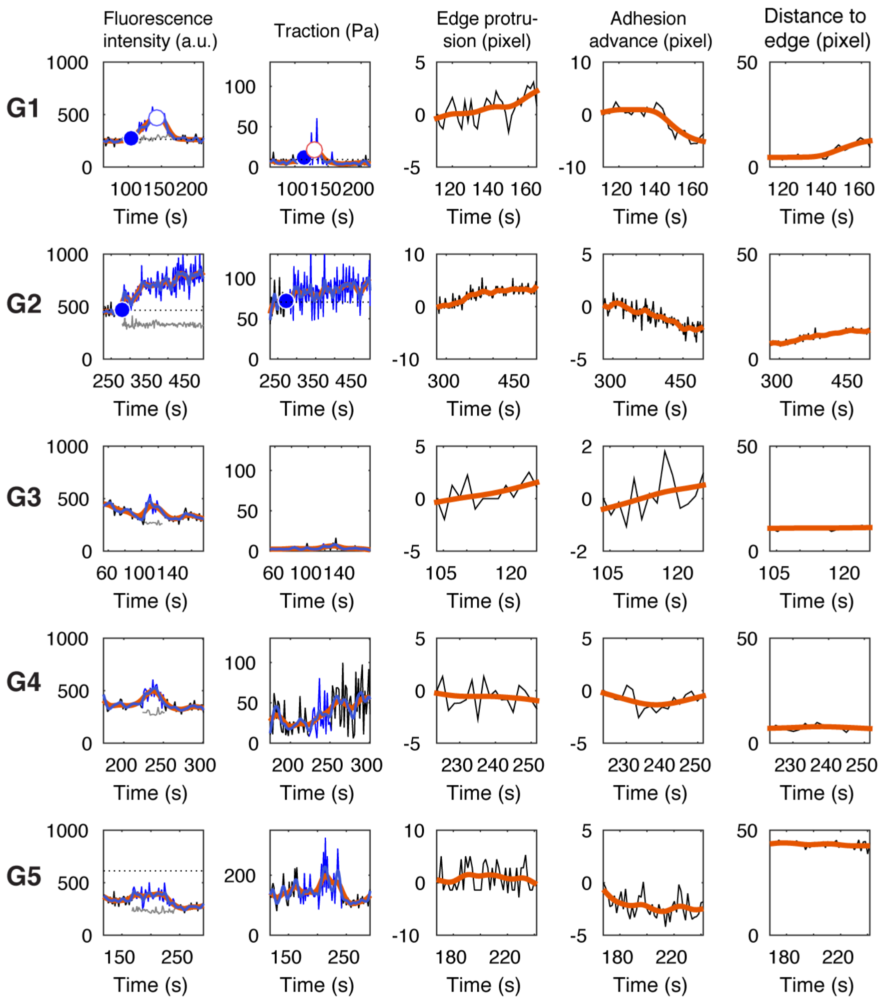

*Supplementary Figure 5. Representative time series of fluorescence intensity, traction magnitude, edge protrusion speed, adhesion sliding speed, and distance to closest the edge, for the five NA groups (G1-G5). The names for all of the features are listed in Supplementary Table 1.*

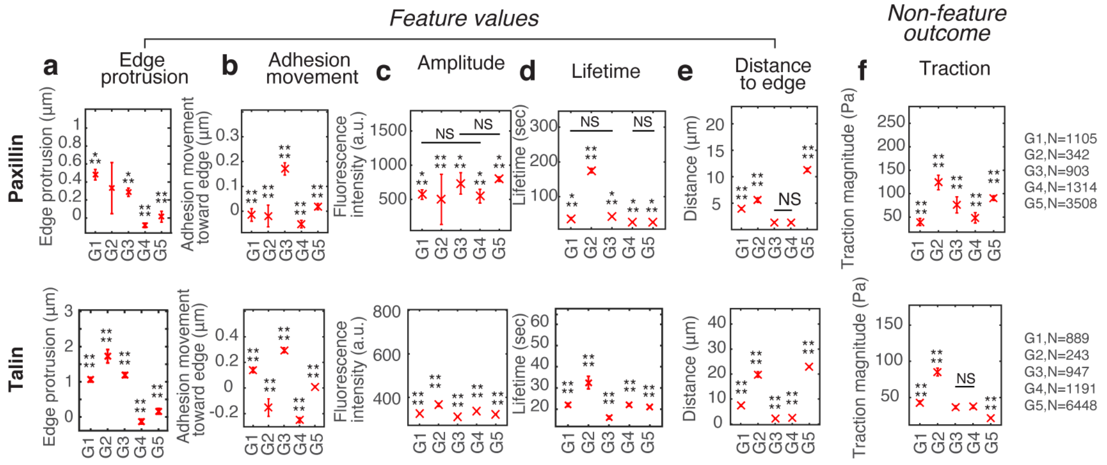

*Supplementary Figure 6. Differences in the five different feature values for all five NA groups, i.e., G1, G2, G3, G4, and G5, as classified with paxillin-mGFP and talin-mGFP. The traction magnitude, which is a non-feature outcome (e.g., is not part of the SVM training set), shows significant differences among classifications groups, similar to differences found from vinculin-mGFP experiments in Fig 1j-n. The sample number per each group is summarized at the right side of each row. Adhesion numbers are extracted from 4 cells for paxillin and 6 cells for talin.*

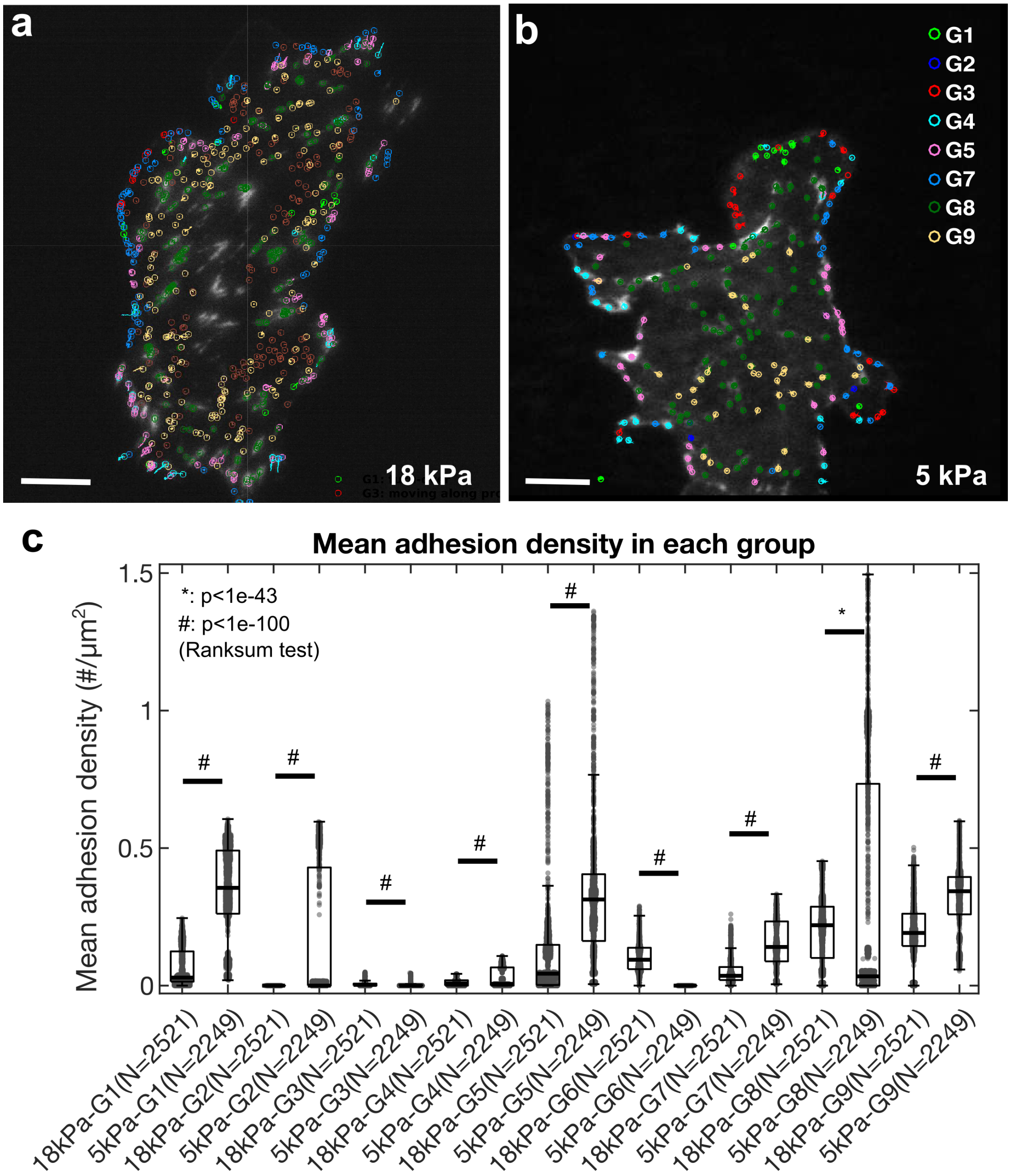

*Supplementary Figure 7. Classification shifts due to substrate stiffness.* ChoK1 cells, transfected with vinculin-GFP, were cultured and imaged on 18 kPa and 5 kPa silicone gel substrates with TIRF. (a) Color-coded classes of kinematically and kinetically different adhesions, tracked and classified using a filter-based classifier (Table S3) for a cell on an 18 kPa and (b) a 5 kPa gel. (c) Mean adhesion density for each adhesion class (G1-G9) as a function of substrate stiffness. N represents the number of frames during an active cell protrusion. Total 9 movies for 18 kPa gel and 6 movies for 6 kPa gel were captured. Note that all classes exhibit significant differences between the two stiffness conditions, particularly in newly assembling NAs, G1 and G2 and large FAs in G8.

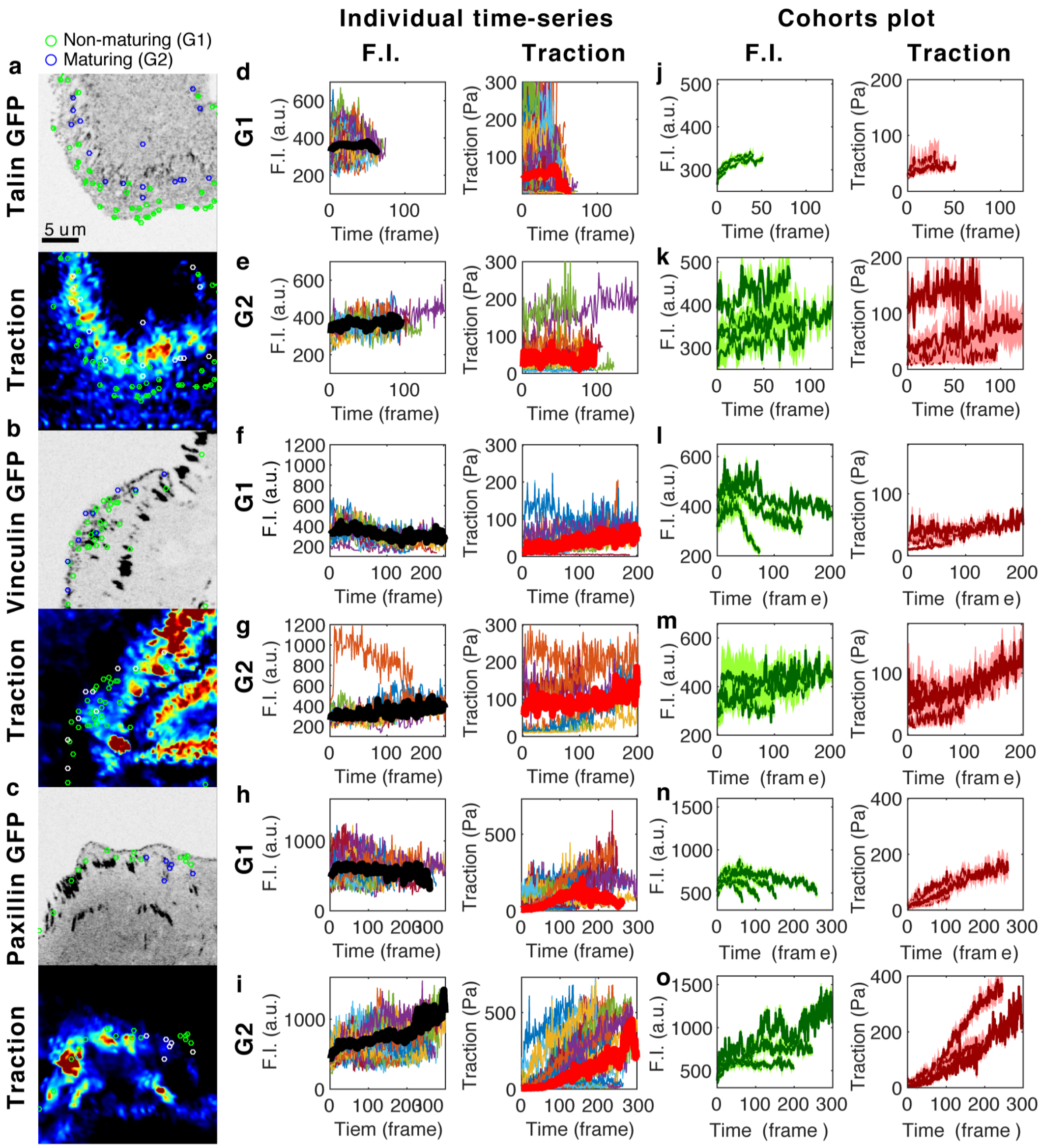

*Supplementary Figure 8. The time-series of fluorescence intensity and traction force within non-maturing (G1) and maturing (G2) NAs for talin, vinculin and paxillin show heterogeneity but "increasing" trend within their lifetimes. Images of GFP-tagged (a) talin, (b) vinculin, and (c) paxillin. Their corresponding traction maps are shown immediately below each image. (d-i) Individual time-series of fluorescence intensity (left column) and traction magnitude (right column) of all non-maturing (d,f,h) and maturing (e,g,i) NAs from the time-lapse images of talin (d-e), vinculin (f-g), and paxillin (h-i). (j-o) Cohorts plots, in which time-series with similar lifetime are grouped and the average time-series per group is plotted with 95% confidence interval, of fluorescence intensity (left column) and traction magnitude (right column) of non-maturing (j,l,n) and maturing (k,m,o) NAs from talin (j-k), vinculin (l-m) and paxillin (n-o).*

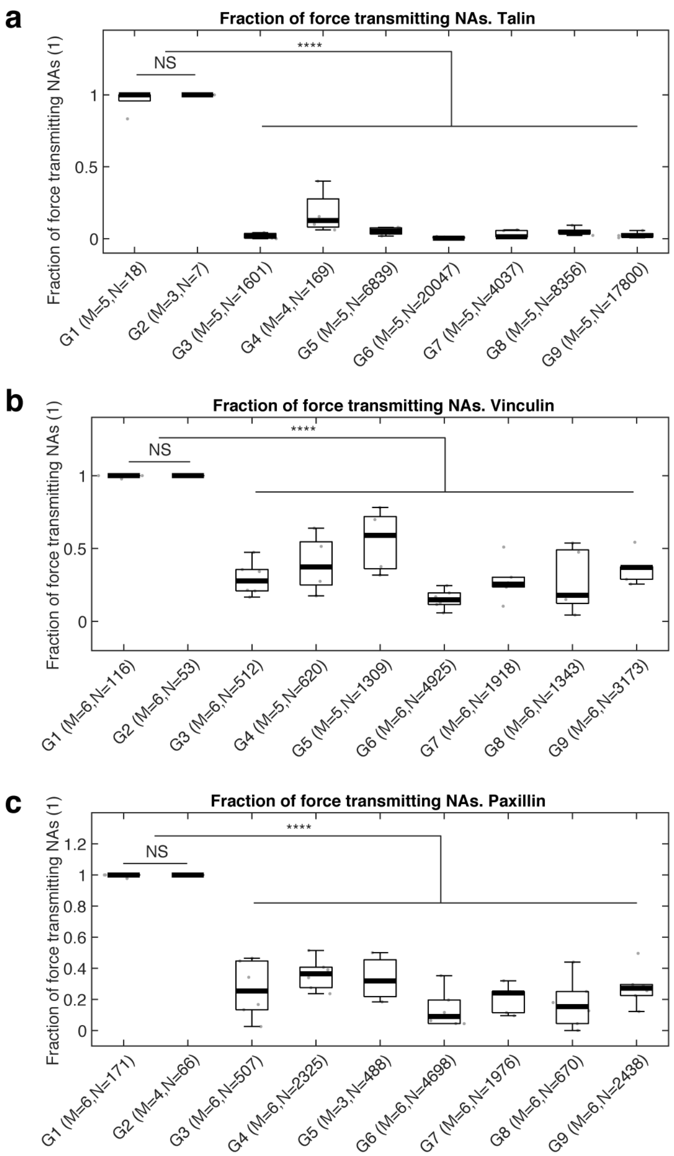

*Supplementary Figure 9. Box plots of fractions of force-transmitting NAs for each classification group for (a) talin, (b) vinculin, and (c) paxillin. Note that nearly 100% of non-maturing (G1) and maturing (G2) NAs generate force, which is much higher than other adhesion classes (G3-G9). Statistical differences among G3-G9 are purposefully not shown.*

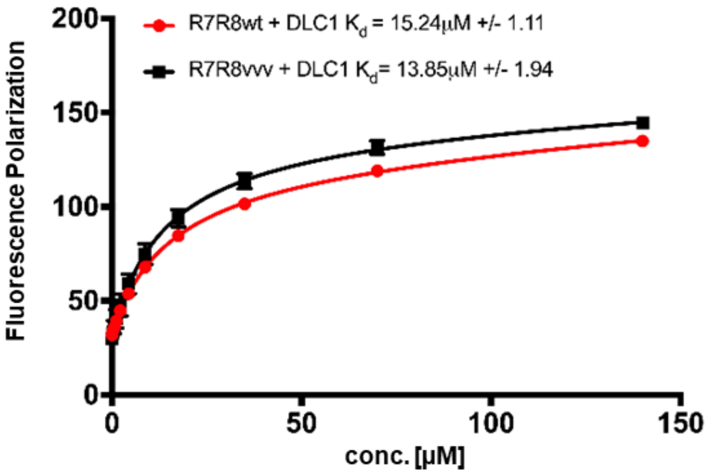

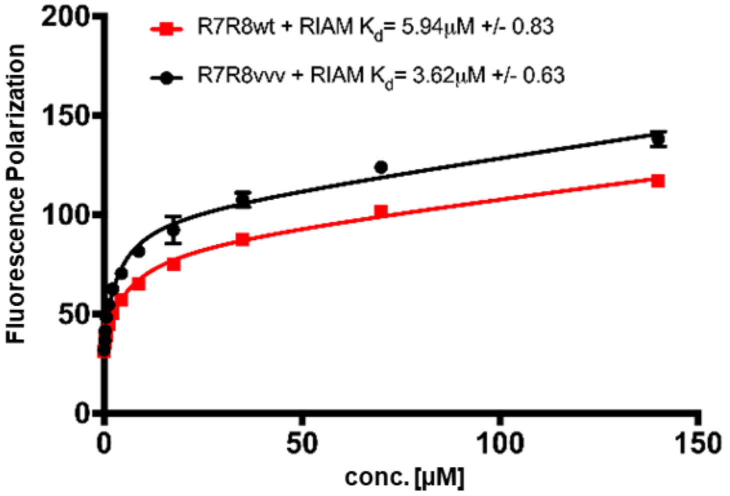

*Supplementary Figure 10. Fluorescence polarization assay showing the binding affinities for R8 ligand peptides from (top) RIAM TBS1 and (bottom) DLC1 with wildtype and R7R8vvv mutants.*

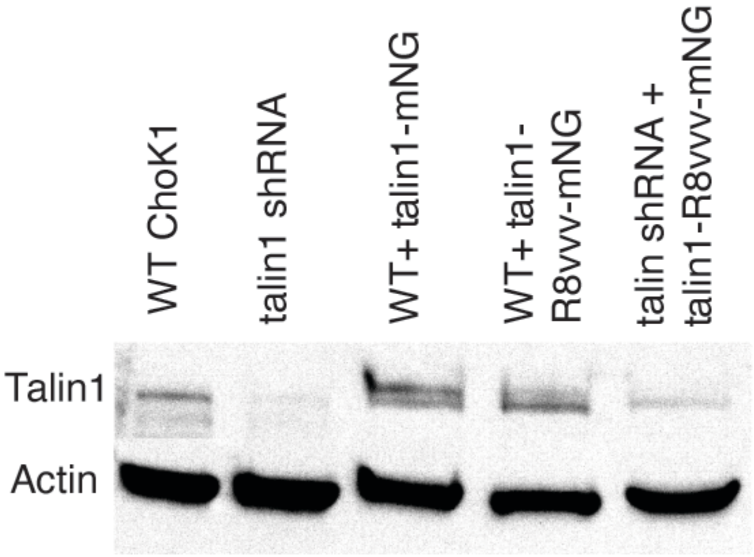

*Supplementary Figure 11. Western blot for talin in wild-type ChoK1 cells, ChoK1 with shRNA-mediated knockdown of talin, wild-type cells with ectopic expression of talin1-mNG, wild-type cells with ectopic expression of talin1-R8vvv-mNG, and ChoK1 cells with shRNA-mediated knockdown of talin and ectopic expression of talin1-R8vvv-mNG. Actin is shown as a loading control.* Note that the double bands in the 3rd and 4th lanes is likely due to the presence of both endogenous and ectopic talin. The presence of a fluorescent protein tag results in a slight shift relative to the endogenously expressed talin. Also note the near-complete shRNA-mediated knockdown of talin expression.

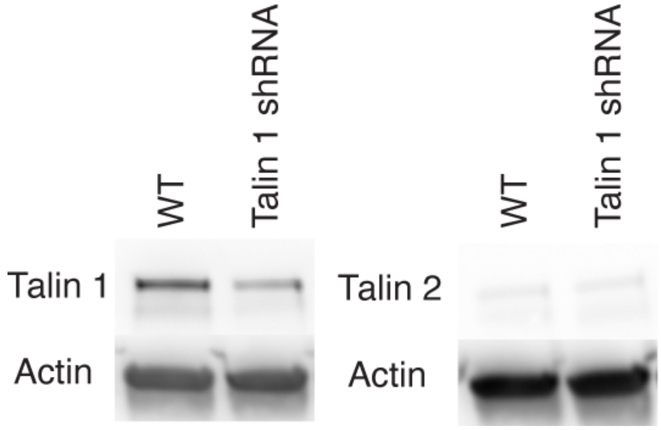

*Supplementary Figure 12. Western blot of talin1 and talin 2 in wild-type and knockdown (shRNA) cells. A blot of actin is shown as a control.*

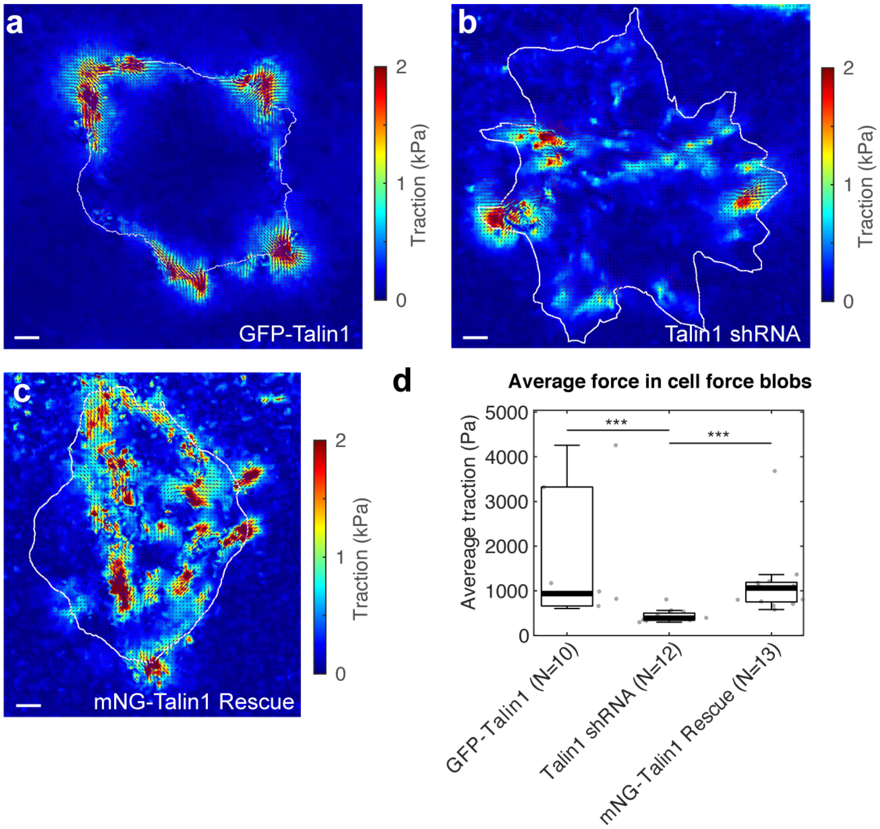

*Supplementary Figure 13. Traction forces generated by ChoK1 cells expressing the talin1 shRNA are smaller than GFP-talin1 and mNG-talin1 rescue cells. (a-c) Traction map with force vectors for ChoK1 celsl with (a) GFP-talin1 overexpression, (b) talin1 shRNA, and (c) mNG-talin1 rescue. (d) The average traction at force blobs, i.e., local forces showing significant force magnitude, for each of the three conditions. Note that average traction is much lower in talin1 shRNA than the two control conditions. ***: $p<1x10^{-3}$ by Mann-Whitney's U-test.*

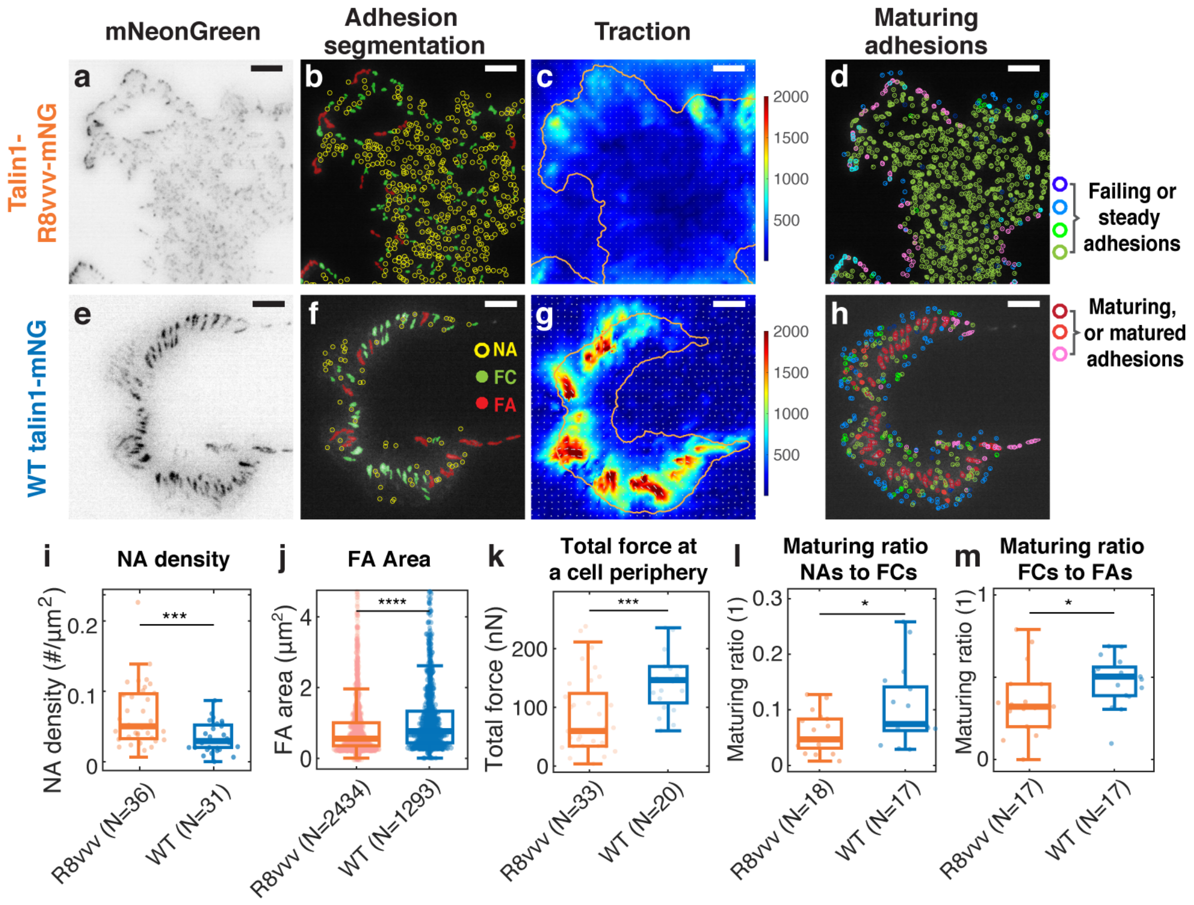

*Supplementary Figure 14. Expression of the talin1 R8vvv mutant results in formation of denser NAs, smaller FAs, lower traction, and less maturing adhesions compared to expression of the talin wildtype. (a-h) Adhesion and traction phenotype of a representative IMCD cell on 5 kPa gel substrates expressing talin R8vvv mutant (a-d) vs WT talin1 (e-h). (a,e) inverted talin-mNeonGreen images. (b,f) detection of NAs, FCs and FAs, color-code of them shown in f. (c,g) traction maps, and (d,h) trajectories of NAs, FCs, and FAs classified via the trained classifier. (i-m) Box plots of NA density (i),FA area (j), total traction integrated over cell's perimeter area (k), the fraction of NAs maturing to FAs relative to all NAs (l) and of the fraction of FCs maturing to FAs (relative to all FCs) (m) for talin1 R8vvv-mNG and WT talin1-mNG rescue, respectively. N in i, k, l, m represents the number of independently imaged cells while N in panel j represents the number of FAs in M=36 and M=31 cells for R8vvv and WT talin1 conditions. Scale bars in panels a-h: 5 µm. *: $p<0.05$, ***: $p<1x10^{-3}$, and ****: $p<1x10^{-10}$ by Mann-Whitney U test.*

### Supplementary Movie Legends

**Supplementary Movie 1**: Time-lapse images of GFP-tagged vinculin in a ChoK1 cell, overlaid with adhesion trajectories and classification state from the support vector machine (SVM)-based machine-learning. Different colors represent different classes: G1 (light green), G2 (dark blue), G3 (red), G4 (light blue), G5 (yellow), G6 (cyan), G7 (pink), G8 (dark green), and G9 (brown). The time interval per frame: 2 seconds. Playing speed: 25 frames/sec. Duration of the movie: 12 minutes. Scale bar: 5 μm.

**Supplementary Movie 2**: Time-lapse images of GFP-tagged talin in a ChoK1 cell. The time interval per frame: 1.644 second. Playing speed: 25 frames/sec. Duration of the movie: 8 minutes 13 seconds. Scale bar: 5 μm.

**Supplementary Movie 3**: Time-lapse images of traction force magnitude for a ChoK1 cell expressing GFP-talin. Traction forces are reconstructed from the bead images using high-resolution, L1-regularized, TFM software (Han et al., 2015). The color scale is the same as one at Fig. 1a, i.e., 0 (blue) – 300 Pa (red). The time interval per frame: 1.644 second. Playing speed: 25 frames/sec. Duration of the movie: 8 minutes 13 seconds. Scale bar: 5 μm.

**Supplementary Movie 4**: Time-lapse images of GFP-tagged talin in a ChoK1 cell, overlaid with adhesion trajectories and classification state from the support vector machine (SVM)-based machine-learning. The color coding is the same as in the legend of Movie S1. The time interval per frame: 1.644 second. Playing speed: 25 frames/sec. Duration of the movie: 8 minutes 13 seconds. Scale bar: 5 μm.

**Supplementary Movie 5**: Time-lapse images of GFP-tagged vinculin in a ChoK1 cell. The time interval per frame: 2 seconds. Playing speed: 25 frames/sec. Duration of the movie: 12 minutes. Scale bar: 5 μm.

**Supplementary Movie 6**: Time-lapse images of traction force magnitude for a ChoK1 cell expressing GFP-vinculin. Traction forces are reconstructed from the bead images using high-resolution, L1-regularized, TFM software (Han et al., 2015). The color scale is the same as one at Fig. 1b, i.e., 0 (blue) – 1800 Pa (red). The time interval per frame: 2 seconds. Playing speed: 25 frames/sec. Duration of the movie: 12 minutes. Scale bar: 5 μm.

**Supplementary Movie 7**: Time-lapse images of GFP-tagged paxillin in a ChoK1 cell. The time interval per frame: 1.644 second. Playing speed: 25 frames/sec. Duration of the movie: 8 minutes 13 seconds. Scale bar: 5 μm.

**Supplementary Movie 8**: Time-lapse images of traction force magnitudes generated by a ChoK1 cell expressing GFP-paxillin. Traction forces are reconstructed from the bead images using high-resolution, L1-regularized, TFM software (Han et al., 2015). The color scale is the same as one at Fig. 1c, i.e., 0 (blue) – 1000 Pa (red). The time interval per frame: 1.644 second. Playing speed: 25 frames/sec. Duration of the movie: 8 minutes 13 seconds. Scale bar: 5 μm.

**Supplementary Movie 9**: Time-lapse images of GFP-tagged paxillin in a ChoK1 cell, overlaid with adhesion trajectories and classification state from the support vector machine (SVM)-based machine-learning. The color coding is the same as in the legend of Movie S1. The time interval per frame: 1.644 second. Playing speed: 25 frames/sec. Duration of the movie: 8 minutes 13 seconds. Scale bar: 5 μm.
